# Exploring the Chain Release Mechanism from an Atypical Apicomplexan Polyketide Synthase

**DOI:** 10.1101/2023.05.23.541938

**Authors:** Aaron M. Keeler, Porter E. Petruzziello, Elizabeth G. Boger, Hannah K. D’Ambrosio, Emily R. Derbyshire

## Abstract

Polyketide synthases (PKSs) are megaenzymes that form chemically diverse polyketides and are found within the genomes of nearly all classes of life. We recently discovered the type I PKS from the apicomplexan parasite *Toxoplasma gondii, Tg*PKS2, which contains a unique putative chain release mechanism that includes ketosynthase (KS) and thioester reductase (TR) domains. Our bioinformatic analysis of the thioester reductase of *Tg*PKS2, *Tg*TR, suggests differences in putative apicomplexan reductase domains compared to other systems and hints at a possibly conserved release mechanism within the apicomplexan subclass Coccidia. To evaluate this release module, we first isolated *Tg*TR and observed that it is capable of 4 electron (4e^-^) reduction of octanoyl-CoA to the primary alcohol, octanol, utilizing NADH as a cofactor. *Tg*TR was also capable of generating octanol in the presence of octanal and NADH, but no reactions were observed when NADPH was supplied as a cofactor. To biochemically characterize the protein, we measured the catalytic efficiency of *Tg*TR using a fluorescence assay and determined the *Tg*TR binding affinity for cofactor and substrates using isothermal titration calorimetry (ITC). We additionally show that *Tg*TR is capable of reducing an acyl carrier protein (ACP)-tethered substrate by liquid chromatography mass spectrometry and determine that *Tg*TR binds to holo-*Tg*ACP4, its predicted cognate ACP, with a *K*_D_ of 5.75 ± 0.77 µM. Finally, our transcriptional analysis shows that *Tg*PKS2 is upregulated ∼4-fold in the parasite’s cyst-forming bradyzoite stage compared to tachyzoites. Our study identifies features that distinguish *Tg*PKS2 from well-characterized systems in bacteria and fungi, and suggests it aids the *T. gondii* cyst stage. Together, this work increases our knowledge of PKS thioester reductase domains and advances our understanding of unconventional polyketide chain termination mechanisms.

## INTRODUCTION

Natural products comprise a large class of bioactive molecules with uses ranging from anti-cancer and anti-parasitic therapeutics to biofuels and insecticides.^1–6^ As such, the biomachinery responsible for these compounds is exceptionally diverse. Polyketides, formed by polyketide synthases (PKSs), comprise the carbon scaffolds of many pharmaceutically important compounds.^1^ Among PKSs, modular type I PKSs are perhaps the most well-characterized. These proteins are megaenzymes that are organized into distinct modules composed of enzyme domains tethered by short amino acid linkers. Each module functions to extend the polyketide chain through acylation of a starter unit by an acyltransferase (AT) domain followed by substrate transacylation onto a cognate acyl carrier protein (ACP) and finally a Claisen-like condensation with a ketosynthase (KS) domain.^7,8^ Additional tailoring enzymes can be present that modify the growing backbone metabolite. At the end of these assembly-line PKSs, substrate release must occur to liberate the completed product which can be further derivatized by post-PKS enzymes.

Multiple chain-release mechanisms are known to exist such as canonical type I and II thioesterases (TEs), oxygenases, FabH-like enzymes, standalone cyclases, and KS or AT-mediated chain release.^9–13^ Of particular importance are thioester reductase domains (TR) which release the completed metabolite through either a two electron (2e^-^) or four electron (4e^-^) reduction of an acyl-thioester to an aldehyde or primary alcohol, respectively.^14^ Given these tail groups, natural products produced by TR domains often have significant biological relevance and activity such as leupeptin (protease inhibitor), myxochelin A (siderophore), and azinomycin B (DNA alkylating agent), and many can undergo further functionalization by post-processing enzymes.^15–17^ However, few TR domains have been biochemically characterized to date, causing a lack of knowledge on their conserved properties, particularly in atypical systems.^18–20^

Structural studies of TR domains have demonstrated that they belong to the superfamily of short-chain dehydrogenase/reductase (SDR) enzymes involving a tyrosine-dependent catalysis mechanism and conserved Rossman fold.^9,14^ TR domains typically consist of a Ser/Thr-Tyr-Lys catalytic triad and their structures can be broken down into two domains. The more conserved N-terminal domain, which harbors a Rossman fold with a glycine repeat, GxxGxxG motif, is responsible for NAD(P)H cofactor binding. The more divergent C-terminal domain, which has a more flexible structure, is involved in substrate and protein binding. Previous structural and biochemical studies have shown that catalysis proceeds through a nonprocessive mechanism, where the substrate and used cofactor must leave the active site after a reduction step.^18,19^ If a second reduction step occurs, substrate and another cofactor must bind before the reaction proceeds. As some TR domains catalyze a two-step reduction to a primary alcohol, while others only catalyze a reduction to an aldehyde, it is difficult to predict the product formed by these domains from primary sequence analysis. Leys and coworkers determined in a related carboxylic acid reductase (CAR) domain that specific C-terminal motions induce backbone reorientations that block the active site in the absence of acyl-thioester substrates. Thus, this movement prevents unwanted additional aldehyde reductions within the active site. They further showed that the movement of a key aspartate residue near the cofactor binding site, facilitated by C-terminal domain dynamics caused by phosphopantetheine (ppant) arm interactions, prevented aldehyde reduction.^21^ However, it remains unclear if this mechanism is specific to CAR domains or more generally conserved. While important studies have investigated this domain in bacterial and fungal systems, to the best of our knowledge there have been no biochemically characterized TR domains from a protist. Apicomplexans, which include the parasites responsible for malaria, toxoplasmosis, and cryptosporidiosis, encode polyketide synthases with predicted TR domains that have yet to be characterized. For example, the apicomplexan parasite *Toxoplasma gondii* encodes two type I PKSs, *Tg*PKS1 and *Tg*PKS2.^22^ We recently elucidated the domain architecture of the 18.8 kb *Tg*PKS2 (**Figure 1A**).^23^ *Tg*PKS2 has an intriguing putative substrate release containing a KS and TR domain, an arrangement that is rarely observed.^24,25^

**Figure 1.**
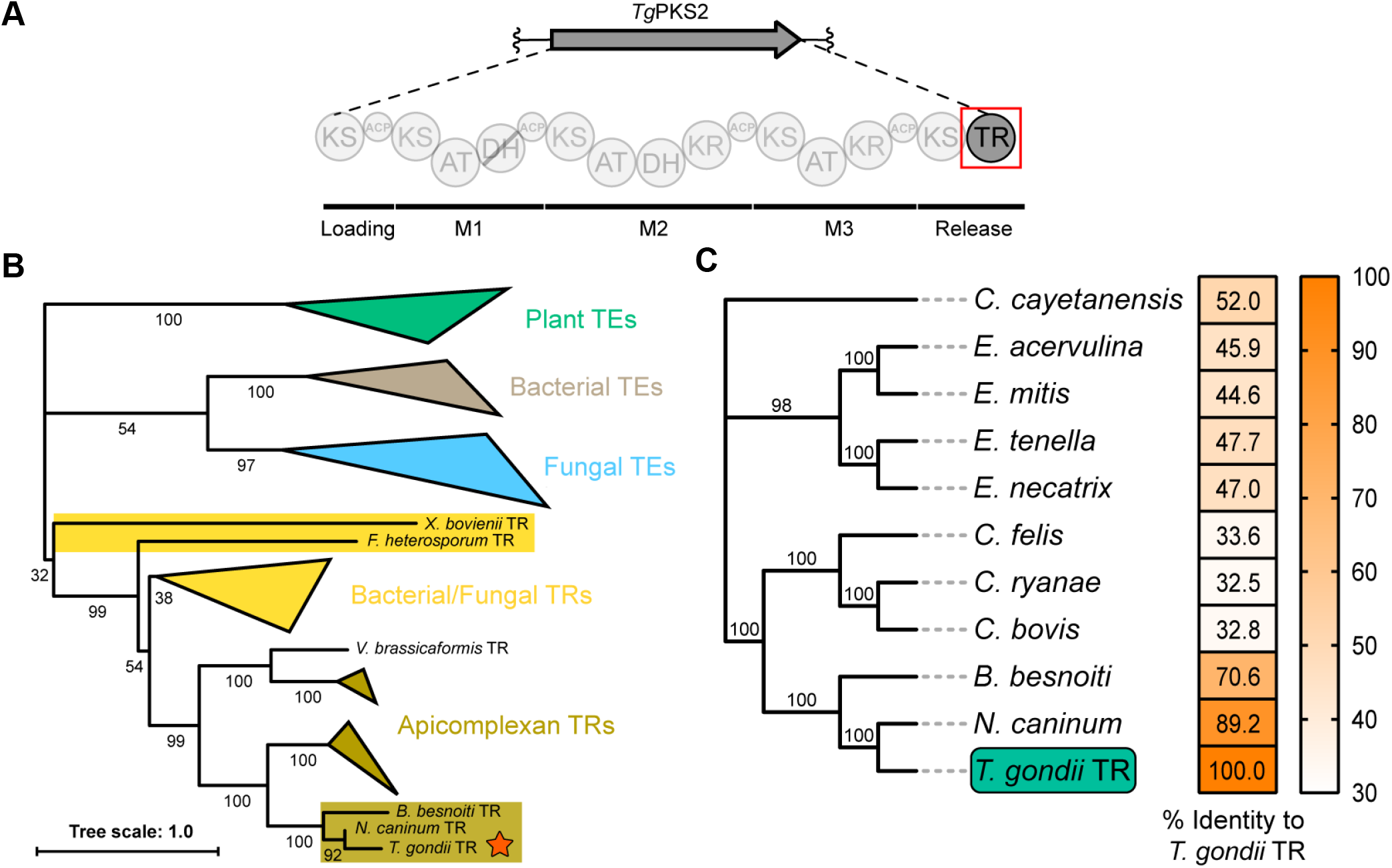
Phylogenetic analysis of TE and TR domains aligned by ClustalW. (A) Domain architecture of *Tg*PKS2 gene cluster. *Tg*PKS2 C-terminus contains a putative thioester reductase (TR) domain that is predicted to release metabolites. (B) Phylogenetic tree of representative TE and TR proteins from known type I and II PKS gene clusters labeled by domain name, *Tg*TR is denoted by a star. Apicomplexan TRs appear to be distinct from previously described systems. (C) Bootstrap consensus tree, using MEGA11, of putative apicomplexan Coccidia TR domains, color denotes percent identity of each domain to *Tg*TR.

Herein, we sought to investigate the *Tg*PKS2 thioester reductase domain, termed *Tg*TR, given the importance of release mechanisms in polyketide formation. Our bioinformatic analysis indicates large differences in putative apicomplexan reductase domains, like *Tg*TR, when compared to better characterized bacterial and fungal systems. *Tg*TR contains an atypical predicted catalytic triad, and we demonstrate that it is capable of 4-electron (4e^-^) reduction *in vitro* of a thioester substrate mimic solely utilizing NADH as a cofactor. We also establish an interaction between *Tg*TR and the predicted cognate ACP, *Tg*ACP4, where *Tg*TR binds to the ACP and reduces an ACP-tethered substrate *in vitro*. We further show that the *Tg*PKS2 gene cluster is upregulated in the bradyzoite stage of *T. gondii* infection, suggesting a role in cysts. Collectively, these findings demonstrate that *Tg*PKS2 is distinct from previously characterized systems in bacteria and fungi and paves the way for further study of PKSs in atypical sources such as apicomplexans and other protists.

## RESULTS

### Bioinformatic Analysis of *Tg*TR

The minimum boundaries of the *Tg*TR domain (5775–6177 amino acids) from *Tg*PKS2 were identified with BLASTP analysis and evaluation of a homology model. We conducted a phylogenetic analysis of *Tg*TR, comparing it to other organisms that encode terminal TR domains. We found that apicomplexan PKS TRs appear to be phylogenetically distinct from those in bacterial and fungal systems (**Figure 1B**). From this we observed that *Tg*TR was most similar to the TR domains from the related apicomplexans *Neospora caninum* and *Besnoitia besnoiti* (∼89% and ∼71%, respectively). With this knowledge, we performed a comparative analysis within the apicomplexan phylum. Although PKS genes are known to exist in many species of apicomplexan, they appear restricted to the Coccidia sub-class.^23,26,27^ These species include *Cryptosporidium* spp., *Eimeria* spp., and *Neospora* spp. among others. Using BLASTP and fungiSMASH we analyzed the best assembled genomes for several species of Coccidia and compared putative protein identity to *Tg*TR (**Figure 1C**). Each prospective TR domain had significant similarity (89–30%) to *Tg*TR.

### Cloning, Expression, and Purification of *Tg*TR

The predicted *Tg*TR domain was analyzed to ensure that the construct contained all previously reported requisite motifs and folds (**Figure 2A**).^28^ This TR construct was cloned from an *E. coli* codon optimized sequence into an expression vector harboring an N-terminal maltose binding protein (MBP) and C-terminal His6-tag. This construct was expressed in *E. coli* BL21(DE3) where we observed the MBP-tag was requisite for the expression of soluble protein. After expression, *Tg*TR was purified by Ni-NTA affinity chromatography followed by anion exchange chromatography to homogeneity (≥95%). Protein purity was determined by SDS-PAGE and identity was confirmed by western blot (**Figure 2B**). Purified *Tg*TR yield was 0.5 mg/L.

**Figure 2.**
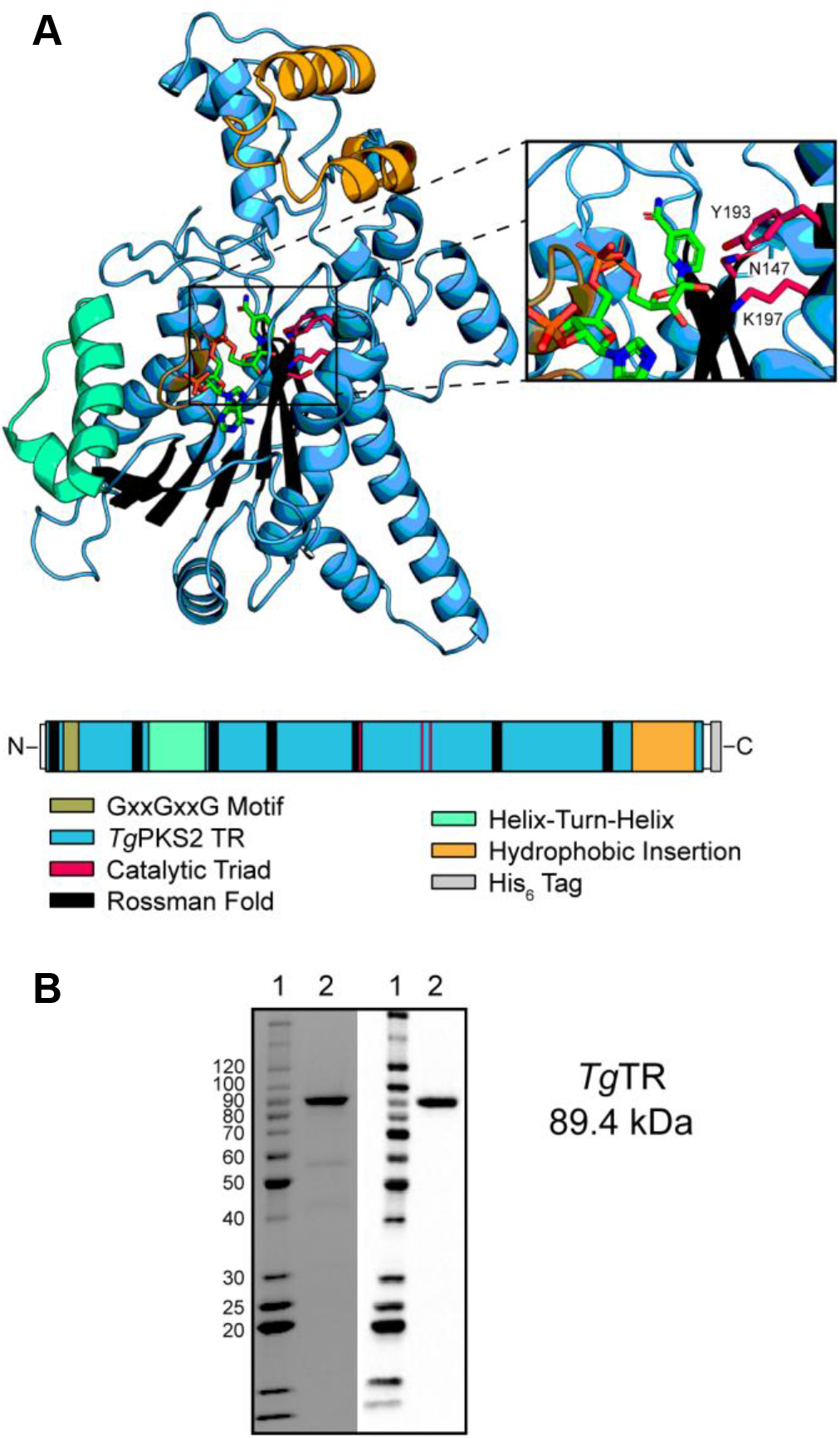
*Tg*PKS2 thioester reductase (TR) domain. (A) Alphafold2 homology model of *Tg*TR (cartoon), active site resolved by alignment of NADPH (sticks, from PDB ID: 4U7W). Inset shows the predicted catalytic triad residues. Annotated motifs and important regions of *Tg*TR are shown below the structure. (B) SDS-PAGE gel (left panel) and western blot (right panel) of purified *Tg*TR. Lane 1, BenchMark ladder; Lane 2, purified *Tg*TR.

### *Tg*TR Utilizes the Cofactor NADH for Reduction

We sought to determine the identity of the *Tg*TR cofactor and probe the ability of the domain to utilize a non-native substrate. To this end, we employed a fluorescence assay using octanoyl-CoA (OCoA) as a substrate analog, as previously reported.^19^ Briefly, reduction of OCoA by *Tg*TR utilizing NADH or NADPH causes cofactor oxidation to NAD+ or NADP+, respectively, which can be continuously monitored by a decrease in fluorescence on a plate reader. Using this assay, we observed consumption of NADH by *Tg*TR (10 µM) in the presence of OCoA substrate, and no activity is seen in a parallel boiled enzyme control reaction (**Figure 3**). Interestingly, a reduction of NADPH was not observed in the presence of *Tg*TR and OCoA. Reactions with *Tg*TR and the corresponding aldehyde, octanal, demonstrated similar reduction of NADH but not NADPH over time (**Figure S1**). Thus, *Tg*TR solely utilizes NADH as a hydride source and appears capable of reducing non-native substrates, OCoA and octanal, indicating some level of promiscuity as has been previously observed for TR domains.^18–20^

**Figure 3.**
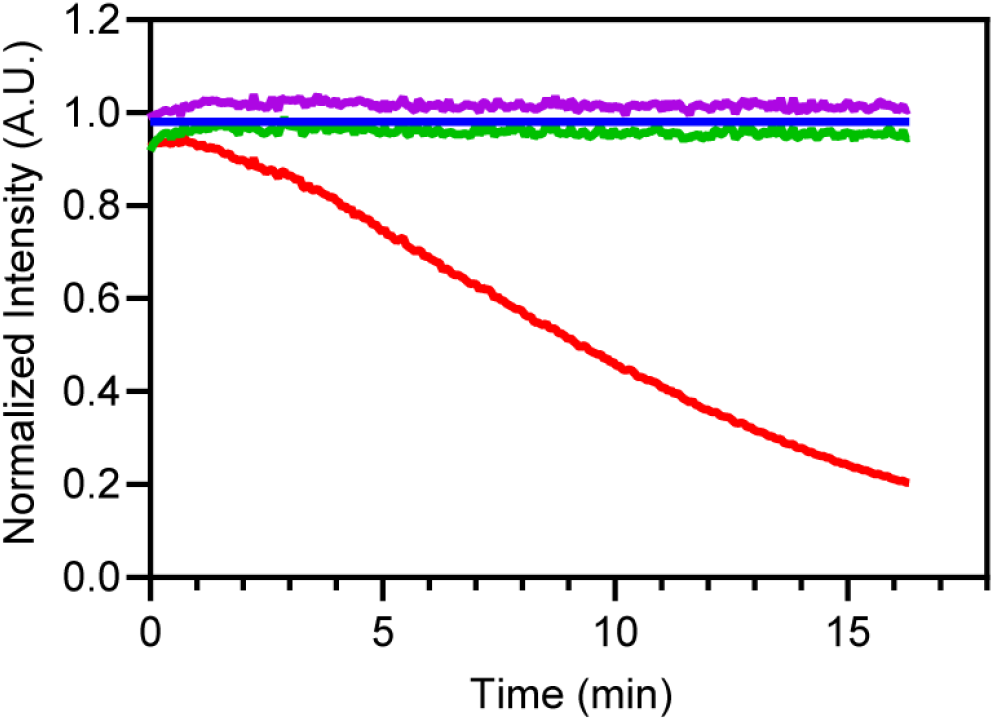
Octanoyl-CoA reduction by *Tg*TR. Normalized fluorescence of NADH control without enzyme (blue trace), NADPH with *Tg*TR (green trace), NADH with *Tg*TR (red trace), and NADH with boiled *Tg*TR (purple trace) monitored 0–16 min. All reactions contained 500 µM octanoyl-CoA and 10 µM *Tg*TR, where indicated. Samples were normalized to the no enzyme control.

### *Tg*TR Catalyzes 4e^-^ Reduction *in vitro*

To validate the reduction of OCoA and determine whether *Tg*TR can perform two rounds of reduction to form the primary alcohol we employed gas chromatography mass spectrometry (GCMS). Purified *Tg*TR (10 µM) was incubated with 500 µM OCoA or octanal for 20 hr at room temperature. Reactions were quenched and extracted with CHCl_3_, derivatized with *N,O*-Bis(trimethylsilyl)acetamide (BSA), and then analyzed by GCMS. We observed that the authentic BSA-derivatized octanol eluted at ∼6.7 minutes with a m/z of 187.20, which corresponds to neutral loss of a methyl group.^19^ In our reaction with *Tg*TR and octanoyl-CoA, a peak with an identical retention time and mass to the octanol standard was observed. This peak was not observed in a boiled *Tg*TR control reaction that was run in parallel (**Figure 4A**).

**Figure 4.**
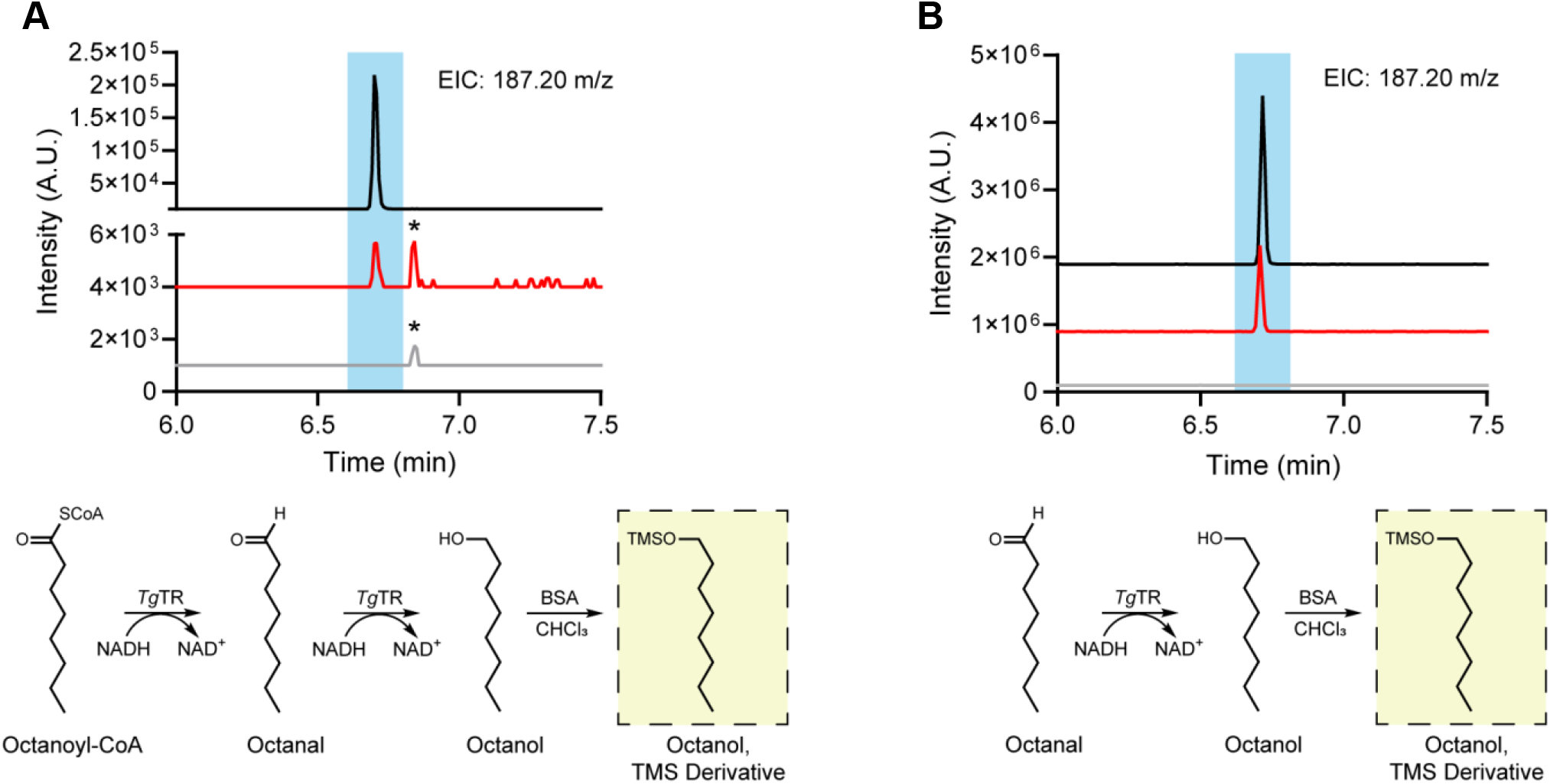
Reduction of thioester and aldehyde substrates by *Tg*TR. (A) GCMS extracted ion chromatogram at 187.20 m/z of the 4 electron reduction of octanoyl-CoA to octanol using NADH, followed by BSA derivatization. An authentic standard of BSA-derived octanol (black trace), BSA-derivatized products with 10 µM *Tg*TR (red trace), and a negative control reaction containing boiled enzyme (grey trace) are shown. * denotes an unidentified impurity present in octanoyl-CoA. (B) GCMS extracted ion chromatogram at 187.20 m/z of the 2 electron reduction of octanal to octanol using NADH, followed by BSA derivatization. An authentic standard of BSA-derived octanol (black trace), BSA-derivatized products with 10 µM *Tg*TR (red trace), and a negative control reaction containing boiled enzyme (grey trace) are shown. Corresponding reactions schemes shown under spectra.

Further, we incubated *Tg*TR with octanal under the same reaction conditions and observed a similar reduction to octanol. Again, this reduction of octanal was not observed in a boiled enzyme control reaction (**Figure 4B**). These experiments indicate that *Tg*TR can perform the 4e-reduction of octanoyl-CoA to octanol and the 2e^-^ reduction of octanal to octanol *in vitro*.

### Kinetic Parameters and Substrate Binding Affinity to *Tg*TR

To further characterize *Tg*TR, purified protein (10 µM) was incubated with varying concentrations of substrate, OCoA or octanal, and the NADH signal was monitored over time. The rate of NADH consumption was determined at each substrate concentration using the aforementioned fluorescence assay and quantitated using a calibration curve with known standards (**Figure S2A**). We confirmed that NADH consumption was linear with respect to time for OCoA and octanal (**Figure S2B and S2C**). Initial burst reaction velocities were plotted as a function of the substrate concentration and subsequently fit to a non-linear regression (**Figure 5A**). Based on these saturation curves, *Tg*TR exhibited an observed *k*_*cat*_ = 0.15 min^-1^ with an observed *k*_*cat*_*/K*_*m*_ = 1.1 ± 0.5 nM^-1^ min^-1^ for octanoyl-CoA. The observed *Tg*TR catalytic efficiency value was slightly improved (∼4-fold) for octanal, where the determined *k*_*cat*_ = 0.64 min^-1^ and *k*_*cat*_*/K*_*m*_ = 4.1 ± 0.6 nM^-1^ min^-1^. While *Tg*TR lacks high sequence similarity (∼24% similarity) to previously characterized *Mycobacterium* spp. TR and CAR domains, their kinetic parameters are comparable.^18,20,21^ To further explore the substrate pocket size we evaluated *Tg*TR activity with the 10-carbon aldehyde, decanal. The determined saturation curves show that *Tg*TR exhibited an observed *k*_*cat*_ = 0.34 min^-1^ with an observed *k*_*cat*_*/K*_*m*_ = 1.5 ± 0.3 nM^-1^ min^-1^ for decanal (**Figure 5B**).

**Figure 5.**
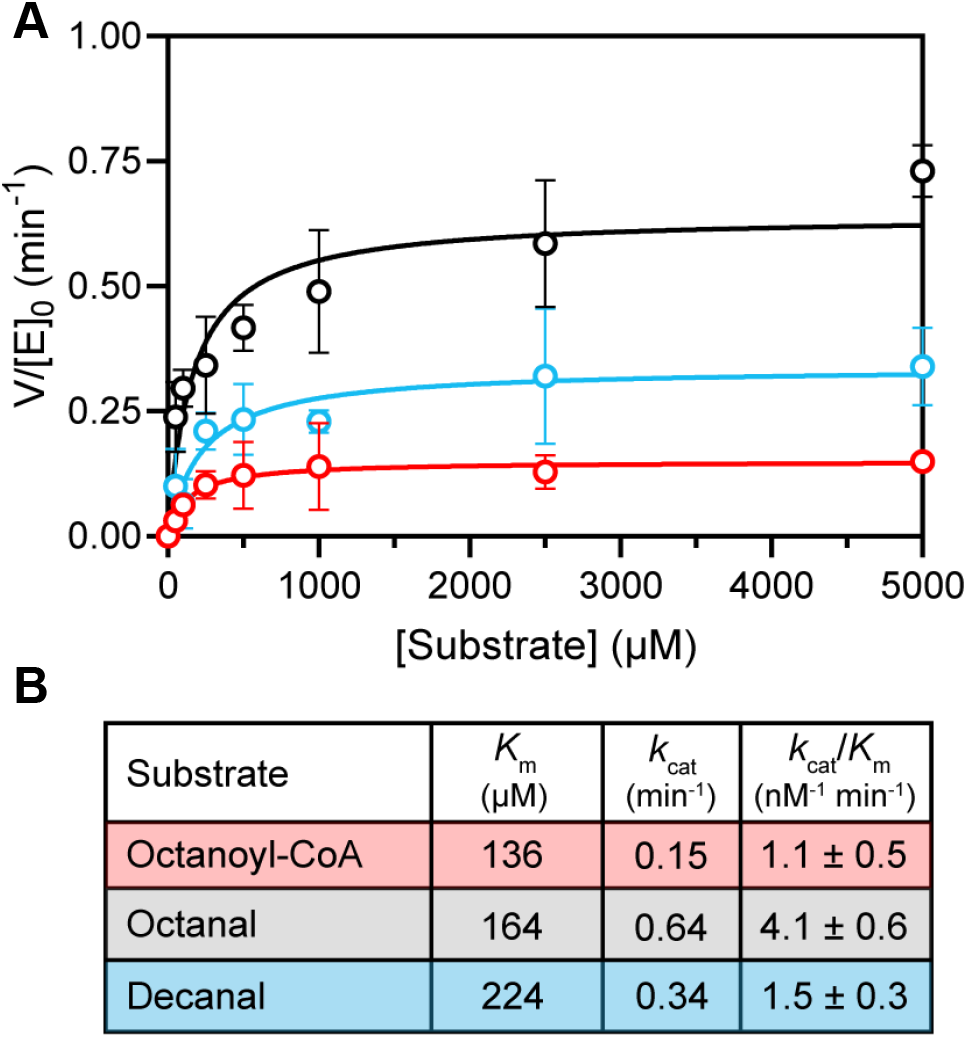
Kinetics of substrate reduction by *Tg*TR. Saturation curve for *Tg*TR in the presence of octanoyl-CoA (red), octanal (black), or decanal (blue). Each reaction contained 10 µM *Tg*TR and 100 µM NADH. (B) Kinetic parameters of substrate reduction. Data are shown as the average ± SEM (n≥3).

Thus far, relatively little is known about the binding affinity of OCoA and octanal, commonly used non-native substrates, to TR domains. Therefore, we first conducted a thermal shift assay (TSA) to determine if a shift in *Tg*TR melting temperature (Tm) occurs upon cofactor or substrate addition. We observed that NADH, octanoyl-CoA, and octanal increased the *Tg*TR Tm compared to the DMSO control, which is indicative of a binding interaction that stabilizes the protein (**Figure S3A and S3B**). We then used isothermal titration calorimetry (ITC) to measure the dissociation constants (*K*_D_) of cofactor and substrates to *Tg*TR. We observed that NADH binds to *Tg*TR with a *K*_D_ of 7.64 ± 0.59 µM, comparable to other TR domains.^18,29^ In contrast, no significant heat change was observed with NADPH, indicating it does not bind to *Tg*TR. Octanoyl-CoA and octanal were found to have *K*_D_ values of 89.6 ± 3.9 µM and 146 ± 31 µM, respectively, when titrated into *Tg*TR in the presence of excess NADH (**Table 1**). Of note, we observed the strongest isotherms in the presence of excess cofactor, consistent with a previous study on ketoreductase domains from *Mycobacteria ulcerans*.^30^

**Table 1.** Binding affinity to *Tg*TR. Binding dissociation (*K*_D_) values for possible *Tg*TR substrates and cofactors determined by ITC. Values shown as average ± SEM (n=2-4 independent measurements). N.O. = not observed.

| Ligand | Cofactor | $K_D$ with *Tg*TR ($\mu\text{M}$) |
| --- | --- | --- |
| NADH | — | $7.64 \pm 0.59$ |
| NADPH | — | N.O. |
| Octanoyl-CoA | NADH | $89.6 \pm 3.9$ |
| Octanal | NADH | $146 \pm 31$ |
| Holo-*Tg*ACP4 | NADH | $5.75 \pm 0.77$ |

### *Tg*TR Binds to the Cognate Acyl Carrier Protein

TR domains do not covalently bind to the growing metabolite directly, as opposed to TE domains; therefore, we sought to evaluate a possible protein-protein interaction between *Tg*TR and the final ACP in *Tg*PKS2, the predicted cognate domain to *Tg*TR, holo-*Tg*ACP4. Using ITC, we observed that holo-*Tg*ACP4 had a *K*_D_ value of 5.75 ± 0.77 µM to *Tg*TR in the presence of excess NADH. This binding affinity is similar to that of ACPs to PKS domains in other systems.^30–33^ ITC data is included in **Table 1 and Figures S4A–E**.

TR domains have been shown to make protein-protein interactions that allow for insertion of the ACP ppant arm into the active site. To better explore this, we modeled this protein-protein interaction first by *Tg*TR electrostatics and then with HAD-DOCK docking software (**Figure 6A**).^34,35^ From this model, the interface area was calculated as ∼1012 Å^2^ using PISA, which is 19.7% of the surface area of *Tg*ACP4 and 5.1% of the surface area of *Tg*TR. This contact area is similar in size to other ACP-complexes and emphasizes the transient nature of these interactions.^32,36–38^ *Tg*ACP4 makes numerous predicted electrostatic interactions with the N- and C-terminal domain of *Tg*TR, particularly via residues within the interhelical region between helix II and III (**Figure 6B**). The serine (S41) residue where attachment of the 4’
s-ppant arm occurs on *Tg*ACP4 is located in reasonable proximity, 12.0 Å, from the center of mass of the predicted catalytic triad: N147, Y193, and K197. This distance is similar to that previously obtained from a model of the myxalamid system.^20^

**Figure 6.**
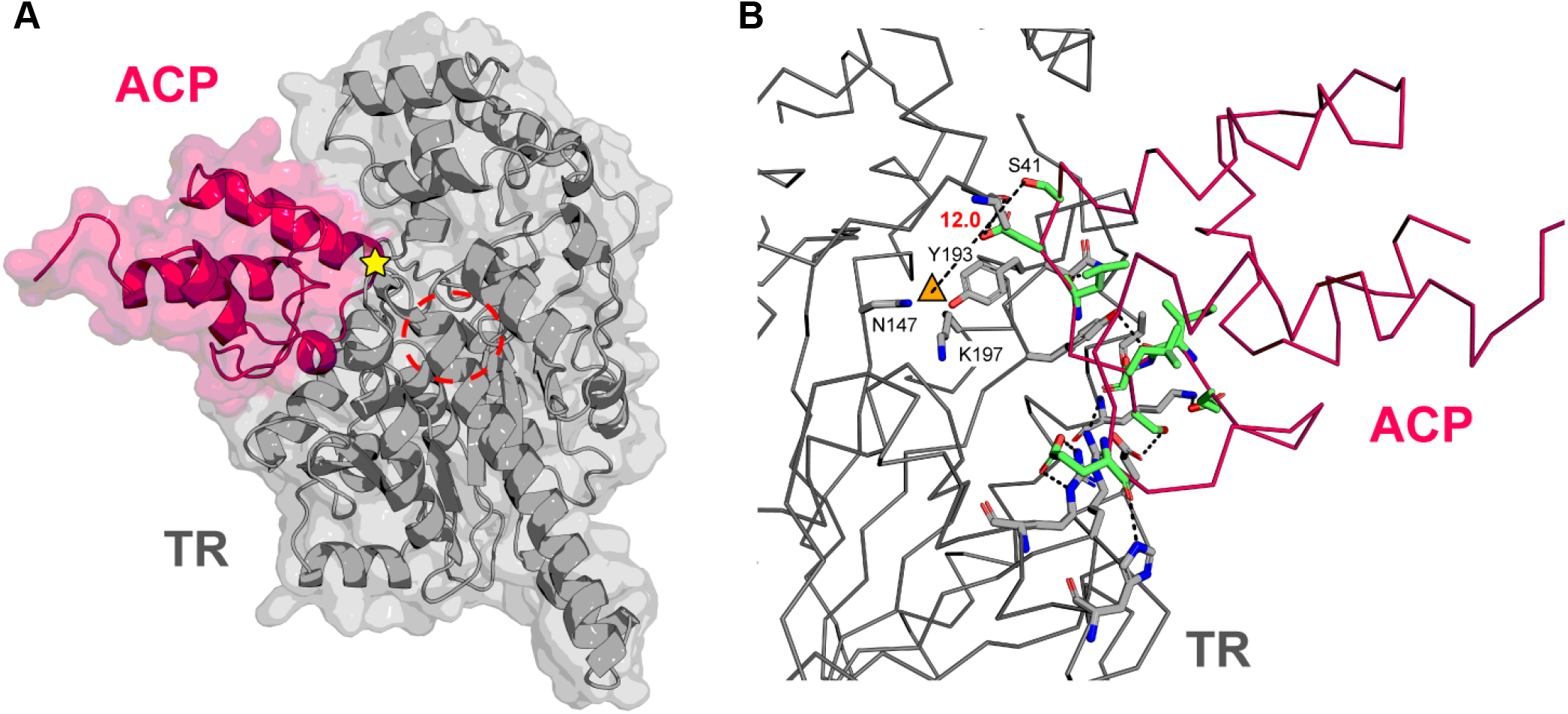
*Tg*TR and *Tg*ACP4 docking interactions. (A) HADDOCK 2.4 protein-protein docking model of *Tg*ACP4 (pink) and *Tg*TR (grey). Location of conserved serine where attachment of 4’ s-ppant arm occurs is denoted by a star, the catalytic triad of *Tg*TR is circled. All analysis was completed on models generated in AlphaFold2. (B) Electrostatic interactions of *Tg*ACP4 residues (green sticks) with *Tg*TR residues (grey sticks) relative to the catalytic triad (orange triangle). Predicted interactions are shown as dotted black lines, proteins are depicted as ribbons. Measurement is shown of *Tg*ACP4 S41 residue where attachment of the 4’-ppant arm occurs to the center of mass (orange triangle) of the predicted catalytic triad: N147, Y193, K197, shown as sticks.

### *Tg*TR Catalyzes Reduction of an ACP-Tethered Substrate

We demonstrated that *Tg*TR can recognize and reduce both a CoA-tethered substrate and free aldehyde. To further characterize this enzyme, we probed whether *Tg*TR can reduce a ppant-tethered substrate attached to the predicted cognate *Tg*ACP4. We incubated purified apo-*Tg*ACP4 (10 µM), with octanoyl-CoA and purified Sfp phosphopantetheinyl transferase to attach the octanoyl moiety (**Figure S5A–C**) at 37 °C for 3 hr. This reaction generated octanoyl-*S*-*Tg*ACP4 (predicted 11,540.1 m/z), which was confirmed by the LCMS ion at 11,540.8 m/z (**Figure S6**). Octanoyl-*S*-*Tg*ACP4 was next incubated with *Tg*TR and NADH overnight at room temperature and the reactions were analyzed by LCMS. We observed a decrease in the peak corresponding to octanoyl-*S*-*Tg*ACP4 upon incubation with *Tg*TR but no apparent decrease of this peak in buffer or a boiled enzyme control reaction. Concurrently, we observed an increase in holo-*Tg*ACP4 upon incubation with *Tg*TR but no apparent increase in buffer or the boiled enzyme control reaction (**Figure 7**). These observations suggest that *Tg*TR can reduce ACP-bound substrates *in vitro*.

**Figure 7.**
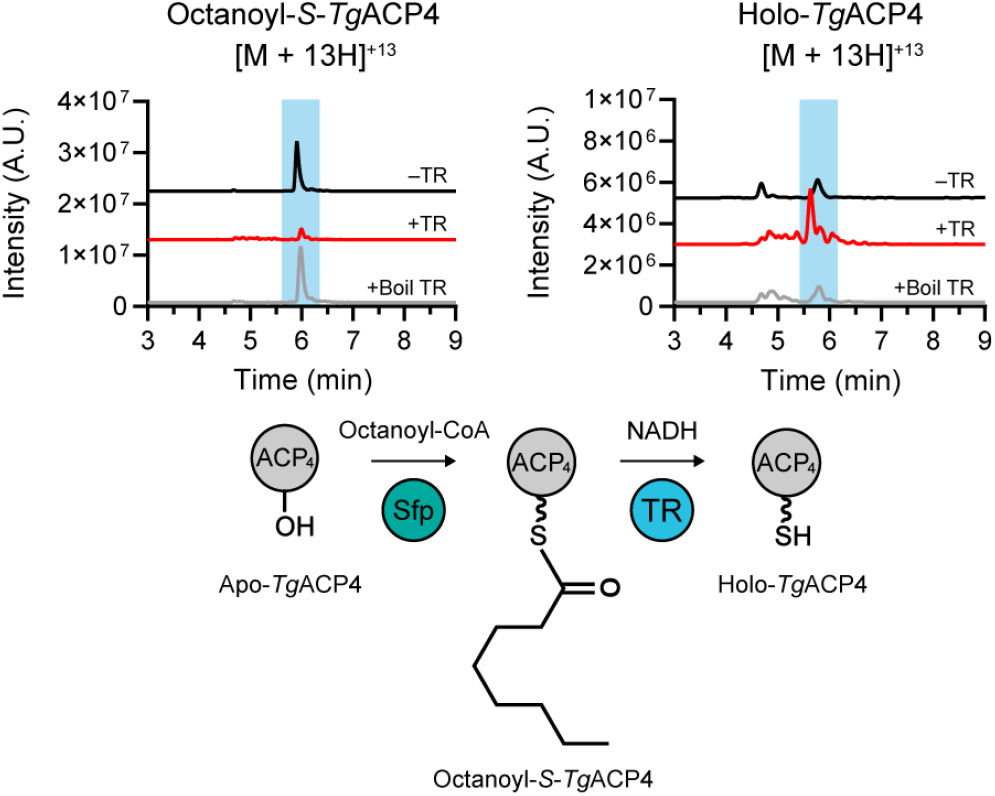
Reduction of ACP-tethered substrate. Representative LCMS extracted ion chromatograms of 13^+^ charge state for octanoyl-*S*-*Tg*ACP4 and holo-*Tg*ACP4 with (red trace) and without (black trace) incubation with *Tg*TR. Reactions with boiled *Tg*TR (grey trace) were included for both substrates. Reaction scheme shown below spectra.

### Probing *Tg*KS5 Activity

Our sequence alignments and homology models indicate that the KS domain proceeding *Tg*TR, *Tg*KS5, contains a conventional Cys-His-His catalytic triad and therefore is predicted to be active. To test this, we examined its capability to perform substrate decarboxylation as previously reported for other loading KS and KS_Q_ domains.^39^ We first cloned, expressed and purified *Tg*KS5 (**Figure S5D**). We also generated malonyl-*S*-*Tg*ACP4 by incubating *Tg*ACP4 with Sfp and malonyl-CoA. *Tg*KS5 was then incubated the malonyl-*S*-*Tg*ACP4 for 3 hr at room temperature and the reaction was evaluated using LCMS. We did not observe formation of acetyl-*S*-*Tg*ACP4 as the LCMS trace with enzyme was the same as that from the boiled enzyme control reaction (**Figure S7**). We additionally incubated *Tg*KS5 with octanoyl-*Tg*ACP4 with and without *Tg*TR, but saw no significant change by LCMS (data not shown).

### Mutational Analysis of *Tg*TR

All previously characterized TR domains from bacteria and fungi contain a highly conserved Thr-Tyr-Lys catalytic triad. Intriguingly, *Tg*TR contains a putative Asn-Tyr-Lys catalytic triad. To investigate how ubiquitous this motif may be, we generated a multiple sequence alignment of *Tg*TR with other characterized TR domains and predicted coccidian TR domains (**Figure S8**). Interestingly, all TR domains retained the critical tyrosine residue, but Coccidia TR domains have amino acid replacements within the catalytic triad. Specifically, the canonical threonine residue was replaced by asparagine or glutamate and the lysine residue was replaced with arginine in some apicomplexan TR domains (**Figure S8 and 8A**). To explore the importance of the catalytic Asn, we generated, expressed, and purified a *Tg*TR N147T mutant to mimic the catalytic triad from bacteria and fungi. We then assessed the ability of *Tg*TR N147T to utilize cofactors. Using a fluorescence assay with octanal substrate, we observed significant consumption of NADH by *Tg*TR N147T (10 µM) and a slight consumption of NADPH (**Figure 8B**). This ability to utilize NADPH was not seen with the wild-type (WT) enzyme.

**Figure 8.**
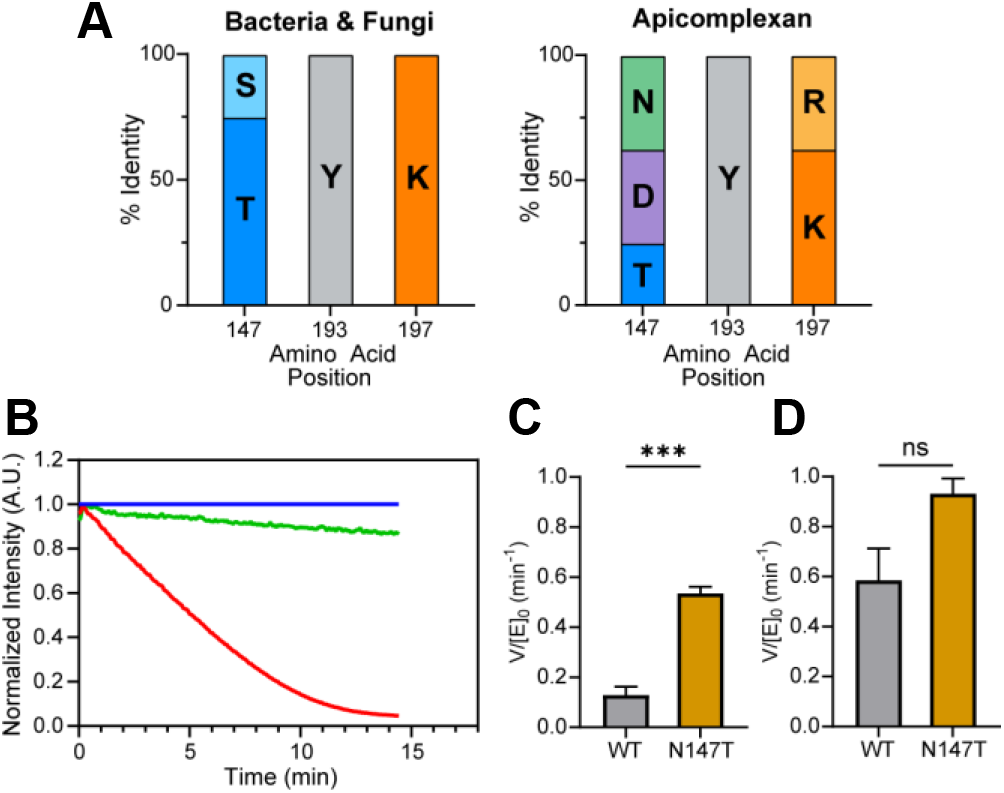
Characterization of *Tg*TR N147T. (A) Catalytic triad identity of 8 representative bacterial/fungal TR sequences (left panel) and 8 representative apicomplexan coccidian TR sequences (right panel). The prevalence of each amino acid (1 letter code) in the sequences is shown as a percentage. (B) Normalized fluorescence of NADH control without enzyme (blue trace), NADPH with *Tg*TR N147T (green trace), NADH with *Tg*TR N147T (red trace). All reactions contained 500 µM octanoyl-CoA and 10 µM *Tg*TR N147T, where indicated. Samples were normalized to the no enzyme control. (C) Relative rates of reduction between WT *Tg*TR and *Tg*TR N147T with saturating octanoyl-CoA (2.5 mM). Data shown as the average ± SEM (n≥3). (D) Relative rates of reduction between WT *Tg*TR and *Tg*TR N147T with saturating octanal (2.5 mM). Data shown as the average ± SEM (n=3). Significance was determined using an unpaired t-test; ***p < 0.001; ns: not significant.

To explore differences in the catalytic activity of the *Tg*TR N147T mutant, we determined single-point binding activity with OCoA and octanal at saturating substrate and NADH concentrations. We observed that *Tg*TR N147T exhibited a significant increase (∼4-fold) in activity with OCoA compared to WT *Tg*TR, but no significant change was observed with octanal (**Figures 8C and 8D**).

### Differential Expression of *Tg*PKS2

*T. gondii* undergoes several different developmental stages during its lifecycle differentiating from the fast-growing tachyzoite into the latent encysted bradyzoite (**Figure S9**). As PKS genes have only been discovered in Coccidia Apicomplexa, which are defined by their ability to generate cysts, we sought to understand if either *PKS1* or *PKS2* gene expression is associated with this life stage using qRT-PCR. *T. gondii* tachyzoites were maintained in continuous culture and then differentiated into bradyzoites by alkaline stress. Three days post-differentiation, mRNA was extracted from both bradyzoite and tachyzoites cultures, cDNA was synthesized, and qRT-PCR using primers targeting *TgPKS1, TgPKS2*, and the bradyzoite stage-specific gene *SAG2D* was performed. As expected, *SAG2D* expression increased in bradyzoite cultures, confirming successful stage transition. *TgPKS2* expression levels increased ∼4-fold in bradyzoite cultures (p=0.03) compared to tachyzoites, while no significant change (p=0.46) was observed in *TgPKS1* (**Figure 9**). This result is consistent with a previous RNA-seq study where a partial gene fragment that we attribute to *TgPKS2* is upregulated 4-fold in bradyzoites when compared to tachyzoites.^40^

**Figure 9.**
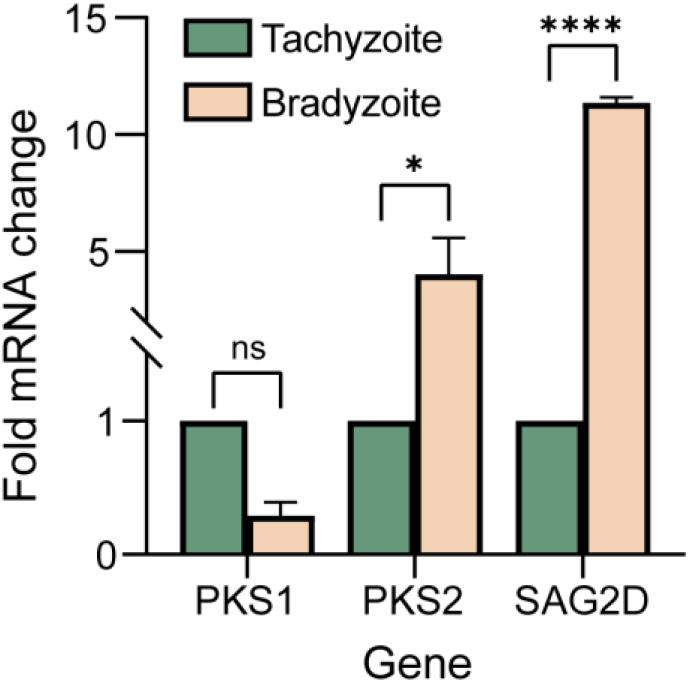
Differential expression of PKS genes in *T. gondii*. Relative transcription levels of *TgPKS1* and *TgPKS2* from tachyzoite or bradyzoite cultures 3 days post-differentiation. The bradyzoite specific gene, *SAG2D*, is included as a marker of stage differentiation. Data shown as the average ± SEM (n=2). Significance was determined using a two-way ANOVA with Sidak’s multiple test comparison of duplicate measurements; ****p < 0.0001; *p<0.05; ns: not significant.

## DISCUSSION

While the presence of PKS biosynthetic gene clusters in apicomplexans has been known for over 20 years, no PKS products have been characterized.^26,27^ This represents a significant gap in our understanding of secondary metabolism in these organisms and the function of molecules they generate. We previously resolved the predicted architecture of *T. gondii* PKS2 and investigated functional domains. The protein contains three defined modules and hypothetical loading and release didomains. Biochemical characterization of the AT and ACP domains revealed unexpected activities.^23,41^ For instance, a unique self-acylation activity is observed with three of the four *Tg*PKS2 ACP domains (*Tg*ACP2–4). These self-acylating ACPs are capable of loading a large scope of CoA substrates onto their ppant arms without an AT domain, which is conventionally required for this activity.^41^ The *Tg*PKS2 putative release di-domain is also unusual as it contains a KS and TR domain. Predominantly a thioester reductase or thioesterase is preceded by an ACP domain. TR domains such as the terminal reductase domain of *Tg*PKS2, appear highly conserved in coccidian parasites. In fact, BLASTP analysis suggests the presence of a KS domain preceding these TR domains is widespread in apicomplexans; however, this is not the case for *Tg*PKS1 which appears to lack a classical release mechanism altogether. Thus, while these domains are distinct from previously characterized systems, our bioinformatic analysis suggests there may be a conserved mechanism for substrate release by PKSs within these organisms. As the release mechanisms in PKSs influence the structure and activity of the final metabolite, we sought to characterize *Tg*TR.

*Tg*TR only utilizes NADH, not NADPH, as the cofactor hydride source for reduction. Interestingly, *Tg*TR N147T, which replaces Asn to mimic the catalytic triad in bacteria, was capable of NADPH reduction, though this activity was limited compared to NADH reduction. These results are interesting as the threonine residue within the catalytic triad is predicted to only play a role in stabilizing the substrate carbonyl through hydrogen bonding, and the asparagine residue would also contain free hydrogen atoms capable of this function. Potentially, this mutation to the smaller amino acid, threonine, opens the binding pocket to accommodate NADPH, or perhaps it repositions a basic residue to stabilize the phosphate moiety. Alternatively, threonine may be positioned to better stabilize the thioester carbonyl as opposed to asparagine, facilitating the initial rate-limiting reduction to the aldehyde. This could account for the increase in the octanoyl-CoA reduction rate in the mutant versus WT *Tg*TR. Our findings suggests that the triad aids in *Tg*TR cofactor selectivity, but biochemical and structural data on TRs with non-typical catalytic residues are critical to establish their selectivity and mechanisms. It remains possible that these proteins exhibit significant structural and mechanistical variability which is often observed within the SDR family of enzymes, or these enzymes might be specifically tuned to accommodate their native substrate scaffolds.^42^

*Tg*TR was able to use octanoyl-CoA, an 8-carbon thioester, as a substrate and catalyze its 4-electron reduction to the primary alcohol, octanol, at a rate comparable to previously characterized reductase enzymes.^18,20,21^ From primary sequence analysis, we were unable to distinguish previously characterized TR domains that undergo 4-electron versus 2-electron reduction of substrates. It is likely that the ability to halt reduction after generation of the aldehyde is more subtle than primary sequence analysis of this domain can discern. For example, there could be structural features such as domain movements during catalysis or intramolecular cyclization like the iminopeptides nostocyclopeptide M1 and koranimine.^21,43,44^ Alternatively, other enzymes may be present that can sequester the aldehyde. In the biosynthesis of cyclizidines, transaminases react with the aldehyde, while oxidases react with the aldehyde in the biosynthesis of tropolone.^45,46^ Thus while *Tg*TR is capable of 4-electron reduction, it is possible that *T. gondii* contains other stand-alone domains that limit reduction to the aldehyde that we have yet to identify.

In addition to octanoyl-CoA, *Tg*TR can reduce free aldehydes (octanal and decanal) as well as react with a CoA-tethered substrate. The determined catalytic efficiency values show a ∼4-fold increase in the reduction rate of octanal compared to octanoyl-CoA, suggesting that the first reduction step is rate-limiting. This is similar to other characterized TR systems and is proposed to stem from the difference in electrophilicity between the thioester and aldehyde.^47^ Interestingly, *Tg*TR binds to octanoyl-CoA with greater affinity than octanal, suggesting interactions between the CoA arm and the reductase binding channel. *Tg*TR can also accommodate a 10-carbon substrate but the catalytic efficiency for decanal was ∼2.7-fold lower when compared to octanal. Although the identity of the native substrate of *Tg*TR is unknown, these findings indicate it can accommodate polyketides between 8–10 carbons in length. Enzymes in this family often exhibit promiscuity, but this substrate length is rational given the modular organization of *Tg*PKS2. Additional studies in *T. gondii* parasites will be critical to definitively determine the native substrate.

Several previously characterized TR domains necessitate the initial binding of the cognate ACP domain to coordinate substrate reduction and are unable to reduce CoA-tethered substrates.^20,21^ While ACP binding is not requisite for *Tg*TR activity, we demonstrated that *Tg*TR binds to *Tg*ACP4 and it is capable of reacting with octanoyl-*S*-*Tg*ACP4. We further sought to understand the role of the KS domain in this reaction as the KSTR di-domain is unusual. In our hands, *Tg*KS5 was inactive *in vitro* using a well-established assay. This suggests that *Tg*KS5 is not capable of a decarboxylative Claisen condensation reaction. This inactivity is not surprising based on its presence at the terminal end of *Tg*PKS2, where decarboxylation of an ACP-tethered substrate would likely be non-productive. It remains possible that *Tg*KS5 is a tailoring enzyme that interacts with the mature *Tg*ACP4-tethered substrate before reductive release by *Tg*TR. In fact, there are several examples of tailoring domains that occur immediately before a release domain, such as methyltransferase domains preceding TR domains in the biosynthesis of stipitatic acid and xenovulene A, or a sulfotransferase domain preceding a thioesterase domain in the curacin A PKS from *Lyngbya majuscula*.^46,48,49^ However, further investigation is necessary to determine the role of this unique di-domain and its influence on the polyketide structure and function.

While the function of the metabolite generated by *Tg*PKS2 is unknown, we show that this gene cluster is transcriptionally upregulated in the bradyzoite stage. Interestingly, *PKS1* is not upregulated during this stage, suggesting they have distinct functions. Among Apicomplexa, PKS gene clusters are restricted to the cyst-forming coccidian sub-class. This association as well as the upregulation of *Tg*PKS2 during the cyst-forming bradyzoite stage, suggests a potential role of the metabolite(s) produced in the cyst lipid wall make up.^22^ In *C. parvum*, a related coccidian parasite, putative PKS and FAS enzymes are upregulated at the lipid-walled oocyst stage, further supporting this proposal^50^, but the structure and function of polyketide specialized metabolites in apicomplexan remains unknown. Currently, treatments with efficacy toward the latent cyst stage of toxoplasmosis are lacking, therefore *Tg*PKS2 or other similar biosynthetic gene clusters in Coccidia may be interesting therapeutic targets.

Altogether, these studies demonstrate the activity of the TR domain in *Tg*PKS2 and deepen our understanding of chain release mechanisms in atypical organisms such as apicomplexan parasites. Our biochemical studies further highlight distinctions in this system from previously characterized PKSs. Further studies on the function of the KS-TR di-domain and the *Tg*TR reduction reaction (2e^-^ vs 4e^-^) in parasites will advance our mechanistic understanding. These may aid in the outstanding questions concerning the *Tg*TR native substrate and the metabolite generated by *Tg*PKS2. Significant challenges exist to answering these questions in the obligate intracellular pathogen *T. gondii*. Further, *Tg*PKS2 is upregulated in the bradyzoite cyst stage, which is technically challenging to work with.^40^ Thus efforts to biochemically characterize domains within this megaprotein are important steps to lay the ground work to elucidate the structure and function of the secondary metabolite product(s) from this apicomplexan PKSs.

## MATERIALS & METHODS

### General

All reagents were purchased from Sigma-Aldrich (St. Louis, MO) unless otherwise indicated. Homology modeling was performed using Deepmind’s Alphafold2 with MMseqs2.^28^ *Tg*TR and *Tg*ACP4 were run separately through the Google Colab notebook, ColabFold: AlphaFold2 w/MMseqs2, with templates and a homooligomeric state of 1.^51^ Protein-protein docking was performed using HADDOCK 2.4.^34,35^ Interface area between *Tg*TR and *Tg*ACP4 was calculated using the PISA server.^52^ All molecular visualizations were generated using PyMOL (Schrödinger).^53^

### Expression and Purification of *Tg*TR

Domain boundaries for the *Tg*TR were determined through fungiSMASH and BLASTP analysis of the *Tg*PKS2 biosynthetic gene cluster against other thioester reductase domains.^24,25^ An Alphafold2 homology model of the proposed domain was evaluated structurally with PyMOL.^53^ *E. coli* codon optimized *Tg*TR was cloned into a pMAL-c2X expression vector (kind gift of Paul Riggs, Addgene #75286) containing an N-terminal maltose binding protein (MBP) tag and a C-terminal His_6_ tag. Plasmids were confirmed by sequencing (Eton Bioscience) and co-transformed into *E. coli* BL21(DE3) with pACYC-GroES/EL-TF (kind gift of Karl Griswold, Addgene #83923) containing GroES/EL and Trigger Factor protein folding chaperones. Cells were plated on LB agar supplemented with 100 µg/mL ampicillin and 25 µg/mL chloramphenicol. For expression, cells were grown in 4 Fernbach flasks containing 1 L of LB media with appropriate antibiotics at 37 °C and 250 rpm. At OD_600_ 1.0, the temperature was reduced to 20 °C and protein expression was induced with 0.1 mM IPTG. After 20 hr at 20 °C cells were collected by centrifugation (4300 g, 30 min) and resuspended in lysis buffer (50 mM KH_2_PO_4_, pH 8.0, 200 mM NaCl, 1 mM benzamidine, 5% glycerol (v/v%), 5 mM β-mercaptoethanol, 10 mM imidazole) supplemented with 1 cOmplete protease inhibitor tablet (Roche). Cells were lysed using a FB120 Sonic Dismembrator (Fisher Scientific) and centrifuged (4300 g, 3 hr). The supernatant was incubated overnight with Ni-NTA resin (Invitrogen) at 4 °C. After stringent washes with buffer containing 10 mM imidazole, bound protein was eluted with a linear imidazole gradient (25–150 mM). Fractions containing *Tg*TR were exchanged into a low salt buffer (25 mM triethanolamine, pH 8.0, 25 mM NaCl, 1 mM DTT, 5% glycerol (v/v%)), applied to an anion exchange HQ/10 column (Applied Biosciences) and eluted with a linear salt gradient. Fractions containing partially purified *Tg*TR were pooled, reapplied to the anion exchange HQ/10 column (Applied Biosciences) and eluted with a linear salt gradient. Protein purity (≥ 95%) was assessed by SDS-PAGE and protein concentration was determined using the Pierce Coomassie Plus Bradford Assay Reagent (Thermo Fisher Scientific). Approximate protein yield was ∼0.5 mg/L for *Tg*TR. Protein was concentrated and stored as 25% glycerol stocks at -80 °C.

### Expression and Purification of Other Protein Constructs

*Tg*ACP4 was cloned into a pET21a(+) expression vector containing an N-terminal T7 tag and a C-terminal His_6_ tag. Sfp was cloned into a pET15b expression vector containing an N-terminal His_6_ tag. *Tg*KS5 was cloned into a pMAL-c2X expression vector (kind gift of Paul Riggs, Addgene #75286) containing an N-terminal maltose binding protein (MBP) tag and a C-terminal His_6_ tag. Plasmids were transformed into *E. coli* BL21(DE3) cells for apo-*Tg*ACP4, Sfp, and *Tg*KS5 or BL21(DE3) BAP1 cells (kind gift of Prof. Chaitan Khosla) for holo-*Tg*ACP4.^54^ After transformation, all cell were plated on LB agar with appropriate antibiotics. For expression, cells were grown in 4 Fernbach flasks containing 1 L of LB media with appropriate antibiotics at 37 °C and 250 rpm. At OD_600_ 0.6, the temperature was reduced to 20 °C and protein expression was induced with 0.5 mM IPTG. After 20 hr at 20 °C cells were collected by centrifugation (4300 g, 30 min) and resuspended in lysis buffer (50 mM KH_2_PO_4_, pH 8.0, 200 mM NaCl, 1 mM benzamidine, 5% glycerol (v/v%), 5 mM β-mercaptoethanol, 10 mM imidazole) supplemented with 1 cOmplete protease inhibitor tablet (Roche). Cells were lysed using a FB120 Sonic Dismembrator (Fisher Scientific) and centrifuged (4300 g, 3 hr). The supernatant was incubated overnight with Ni-NTA resin (Invitrogen) at 4 °C. After stringent washes with buffer containing 10 mM imidazole, bound protein was eluted with a linear imidazole gradient (25–150 mM). Fractions containing partially purified protein were exchanged into a low salt buffer (25 mM triethanolamine, pH 8.0, 25 mM NaCl, 1 mM DTT, 5% glycerol (v/v%)), applied to an anion exchange HQ/10 column (Applied Biosciences) and eluted with a linear salt gradient. Protein purity (≥ 95%) was assessed by SDS-PAGE, and protein concentration was determined using the Pierce Coomassie Plus Bradford Assay Reagent (Thermo Fisher Scientific). Approximate protein yield was 0.5 mg/L for *Tg*ACP4 (apo and holo), 2 mg/L for Sfp, and 0.5 mg/L for *Tg*KS5. Protein was concentrated and stored as 25% glycerol stocks at -80 °C. Amino acid sequences for all protein constructs are shown in **Supplemental Table S1**. SDS-PAGE gels of purified constructs are shown in **Figure S5**.

### Monitoring NAD(P)H Consumption by *Tg*TR

Analysis of NAD(P)H consumption was performed as described previously with minor differences.^18,19^ Briefly, 500 µM octanoyl-CoA (Cayman Chemical), was incubated with 100 µM NADH or NADPH in assay buffer (10 mM Tris-HCl, pH 8.0, 50 mM NaCl, 5% glycerol (v/v%)). Reactions were initiated by addition of 10 µM *Tg*TR to a final volume of 25 µL in a black polystyrene costar 384-well plate (Corning). Fluorescence signals were measured at 460 nm with excitation at 360 nm on an Envision Multimode Plate Reader (PerkinElmer). An identical reaction containing boiled *Tg*TR was used as a negative control. All samples were normalized to NADH control. Data analysis was performed using Prism GraphPad.

### GCMS Analysis of Octanoyl-CoA and Octanal Reduction by *Tg*TR

The reduction of octanoyl-CoA and octanal by *Tg*TR was analyzed using GCMS as similarly described elsewhere.^19^ Briefly, 500 µM octanoyl-CoA or octanal was incubated with 200 µM NADH and 10 µM *Tg*TR in assay buffer (10 mM Tris-HCl, pH 8.0, 50 mM NaCl, 5% glycerol (v/v%)) to a final volume of 500 µL and incubated at room temperature for 20 hr. After incubation, 500 µL CHCl_3_ was added to extract the reaction products and the reactions were vortexed and centrifuged briefly. The CHCl_3_ layer was extracted, dried with Na_2_SO_4_, and filtered through cotton wool. 10 µM *N,O*-Bis(trimethylsilyl)acetamide (BSA) was added to derivatize the reaction products and the samples were analyzed by GCMS (GCMS-QP2010, Shimadzu). Samples were injected (1 µL) through a HP-5MS Ui column (Agilent) using helium as the carrier gas. Temperature was held constant at 70 °C for 2 min and then ramped to 225 °C at a rate of 20 °C/min, the temperature was then held at 225 °C for 2.5 min. An identical reaction containing boiled *Tg*TR was used as a negative control. A BSA-derivatized authentic standard of octanol was used as a positive control comparison.

### *Tg*TR Fluorescence Kinetic Assays

Activity of *Tg*TR was assessed as previously described, with slight modifications.^18,21^ Briefly, reactions were run with 10 µM purified recombinant *Tg*TR with varying octanoyl-CoA, octanal, or decanal concentrations (0–5000 µM) and 100 µM NADH. A control reaction without substrate was run in parallel. All assay components were added to a black polystyrene costar 384-well plate (Corning) to a final volume of 25 µL in assay buffer (10 mM Tris-HCl, pH 8.0, 50 mM NaCl, 5% glycerol (v/v%)). Fluorescence signals were measured at 460 nm with excitation at 360 nm on an Envision Multimode Plate Reader (PerkinElmer). Measurements were taken over a period of 15–20 min and background NADH consumption was subtracted at each timepoint. Data were fit to a nonlinear equation using GraphPad Prism. A calibration curve was completed with known NADH standards (**Figure S2A**).

### Substrate Binding to *Tg*TR by Thermal Shift Assay (TSA)

Protein thermal shift assays were performed by incubating purified *Tg*TR (3 µM) into assay buffer (10 mM Tris-HCl, pH 8.0, 50 mM NaCl, 5% glycerol (v/v%)) and dispensed into white 96-well plates (Roche). Compound (1 mM) or DMSO was added to each well followed by SYPRO orange dye, 5X (Thermo Fisher Scientific) to a final volume of 15 µL. Plates were sealed with clear foil, briefly shaken and centrifuged (1100 rpm x 2 min). Plates were then analyzed using a LightCycler 480 Instrument II (Roche), from 20 °C – 85 °C with a 0.06 (°C/s) ramp rate, 10 acquisitions per °C. Melting temperatures (Tm) were determined by taking the first derivative of each thermal profile using Prism GraphPad.

### Substrate Binding to *Tg*TR by Isothermal Titration Calorimetry (ITC)

Isothermal titration calorimetry (ITC) was utilized to determine the binding affinity of each substrate to *Tg*TR. ITC experiments were carried out using a MicroCal PEAQ-ITC (Malvern Panalytical). The reference cell (300 µL) was filled with water for all experiments. *Tg*TR and all ligands and cofactors were dialyzed into the same buffer (25 mM Tris-HCl, pH 7.6, 25 mM NaCl). A first injection of 0.4 uL was not used in data analysis and was followed by 18 injections of 2 uL, using a stirring speed of 750 rpm, and delay time between injections of either 150 or 400 seconds. Control titrations of ligand into buffer were performed for each substrate. Initially, very small heats (<0.1 µCal/sec per injection) were observed with titration of substrates, preventing isotherm fitting. However, when *Tg*TR was incubated with excess NADH (1 mM) for 20 minutes and then loaded into the sample cell and titrated with ligand, significant heats were observed that could then be fit. This suggests that the interaction with injectant and ligand is isothermic under these experimental conditions without cofactor, a phenomenon that is similar to what was observed in a related system.^30^ Therefore, the sample cell (300 µL) was filled with 10 µM *Tg*TR and 1 mM NADH cofactor (where noted) and the syringe (70 µL) was filled with substrate; NAD(P)H at 400 µM, octanoyl-CoA at 2000 µM, octanal at 2000 µM, or holo-*Tg*ACP4 at 400 µM. Data analysis was performed with MicroCal PEAQ-ITC Analysis Software (Malvern Panalytical) using a single binding site model and visualized using Prism GraphPad. For all isotherm fitting, the stoichiometry, N, was fixed at 1.

### *In vitro* Analysis of Release Domain by LCMS

Sfp was utilized to form malonyl-*S*-*Tg*ACP4 and octanoyl-*S*-*Tg*ACP4 as described similarly.^20,39^ Briefly, 10 µM purified apo-*Tg*ACP4 was incubated with 2 µM Sfp, 1 mM malonyl-CoA or octanoyl-CoA, and reaction buffer (50 mM HEPES, pH 7.4, 12.5 mM MgCl_2_, 1 mM DTT) to a total volume of 250 µL. Reactions were incubated for 3 hr at 37 °C. Afterward, 10 µM purified *Tg*TR and 100 µM NADH or 10 µM purified *Tg*KS5 was added, and the reaction was incubated for an additional 3 hr at room temperature. Samples were then analyzed by LCMS (Agilent 6460 Triple Quadrupole) equipped with a diode array, using a Phenomenex Kinetex EVO C18 100 Å column (2.6 µm, 100 x 3 mm). The mobile phases used were as follows: 100:3 (v:v) H_2_O:MeOH with 0.3% formic acid (A) and 100:3 (v:v) MeCN:H_2_O with 0.3% formic acid (B) with a flow rate of 0.5 mL/min, with the following method: 0–0.5 minutes, 100% A; 0.5–8 minutes, linear gradient from 0–100% B; 8–9 minutes, 100% B; 9–15 minutes, 100% A. All MS data was collected in positive ion mode. MS data was analyzed by MassHunter Qualitative Analysis software (Agilent).

### Phylogenetic Tree Construction

A maximum likelihood phylogenetic tree of TE/TR domain protein sequences from various organisms were compiled, including 4 plant FASII TE sequences, 4 bacterial TE sequences, 5 fungal TE sequences, 7 bacterial/fungal TR sequences, 1 chromerida putative PKS TR sequence, and 11 apicomplexan putative PKS sequences. All sequences used are listed in **Supplemental Table S2**, TR/TE regions were determined by antiSMASH or fungiSMASH and BLASTP.^24,25^ Sequences were aligned using MEGA11 by the ClustalW algorithm and used to construct a neighbor-joining tree using the bootstrap method with 1000 replicates.^55,56^ The bootstrap consensus tree was subsequently visualized in iTol.56,57

### Site-Directed Mutagenesis of *Tg*TR

*Tg*TR N147 was mutated to threonine using the Q5 Site-Directed Mutagenesis Kit (NEB) per the manufacture’s protocol. Primers are included in **Supplemental Table S3**. The *Tg*TR N147T mutant was verified by sequencing (Eton Bioscience) and protein was purified as described above.

### *T. gondii* Growth, RNA Extraction and cDNA Synthesis

*Toxoplasma gondii* type I strain RH tachyzoite cultures, a kind gift from Prof. Sebastian Lourido; were propagated in Vero cells (ATCC, CCL-81) as described previously in a standard tissue culture incubator at 37 °C.^22,39^ Parasites were grown for 72 hr in Dulbecco’s Modified Eagle Medium (DMEM) (Thermo Fisher Scientific) supplemented with 10% heat-inactivated fetal bovine serum (HI-FBS) (v:v), 1% antibiotic-antimycotic (Thermo Fisher Scientific). and subsequently released from Vero cell monolayers by passage through a 20-gauge needle and filtered through a 3 μm polycarbonate membrane (Thermo Fisher Scientific). Parasites were collected by centrifugation (4000 xg, 10 min) and mRNA was extracted using a Quick-RNA Miniprep kit (NEB) according to the manufacturer’s protocol. mRNA (500 ng) was mixed with 1 μL of random primer solution (Thermo Fisher Scientific), to a final volume of 5 μL, heated to 70 °C for 5 min, and then incubated on ice for 5 min. cDNA was synthesized from each reaction mixture using the GoScript Reverse Transcription System (Promega) per the manufacturer’s protocol.

### *T. gondii* Bradyzoite Differentiation

Vero cells were seeded to 90-100% confluency 1 day prior to infection with *T. gondii* tachyzoites in T-150 cm^2^ cell culture flasks (Corning) in DMEM supplemented with 10% HI-FBS (v:v), 1% antibiotic-antimycotic (Thermo Fisher Scientific). Host cells were infected with 50 million *T. gondii* tachyzoites and grown for 6 hours in a standard tissue culture incubator (37°C, 5% CO_2_). After 6-hours, media was removed from the infected flasks, and replaced with RPMI-1640, supplemented with L-glutamine (Gibco), 10% HI-FBS (v:v), 1% antibiotic-antimycotic (Thermo Fisher Scientific) and buffered with 50 mM HEPES adjusted to pH 8.1 with NaOH. To allow differentiation to occur, these cultures were grown at 37°C with no CO_2_ injection, to maintain an alkaline pH for 72 hours.

### qRT-PCR Analysis of Differential Gene Expression

RT-PCR was completed using Taq DNA polymerase (Taq PCR kit, NEB), according to the manufacturer’s protocol. Relative abundance of *T. gondii* bradyzoite-specific gene *SAG2D* (TGGT1_207150), *TgPKS1*, and *TgPKS2* were quantified using SYBR Green I Master reagents with a LightCycler 480 Instrument II (Roche). *T. gondii SAG2D, TgPKS1*, and *TgPKS2* expression in the bradyzoite cultures was compared to the tachyzoite cultures and normalized to the *T. gondii GAPDH* (TGGT1_289690) housekeeping gene. RNA was isolated from 2 biological replicates and qRT-PCR analysis was run in technical triplicates for all samples. All primers used are listed in **Supplemental Table S4**.

## ASSOCIATED CONTENT

Includes supplementary tables, supplementary figures, amino acid sequences, phylogenetic tree sequences, and supplementary results.

This material is available free of charge via the Internet at http://pubs.acs.org.

## AUTHOR INFORMATION

### Author Contributions

A.M.K, P.E.P, E.G.B, and H.K.D performed all experiments. A.M.K and E.R.D wrote the manuscript. All authors edited the manuscript and have given approval to the final version.

### Notes

The authors declare no competing financial interest.

## ACKNOWLEDGMENT

We thank C. Khosla (Stanford University) for providing *E. coli* BL21(DE3) BAP1 cells. We thank P. Silinski (Duke University) for ITC, GCMS, and LCMS assistance. We thank J. Ganley for helpful discussion. We thank the members of the Derbyshire lab for reviewing the manuscript.

## ABBREVIATIONS

PKS: polyketide synthase
TE: thioesterase
TR: thioester reductase
ACP: acyl carrier protein
KS: ketosynthase
ppant: phosphopantetheinyl
mCoA: malonyl-CoA
OCoA: octanoyl-CoA
DTT: dithiothreitol
MeOH: methanol
MeCN: acetonitrile

