## Supplemental Information for "Exploring the Chain Release Mechanism from an Atypical Apicomplexan Polyketide Synthase"

**Table of Contents**

**A. Supplemental Tables**

**Table S1.** **Amino acid and nucleotide sequences for all plasmid constructs.**

**Table S2. Domain name and amino acid sequence of TE/TR domains for phylogenetic trees.**

**Table S3. *Tg*TR site-directed mutagenesis primers.**

**Table S4. qRT-PCR primers.**

**B. Supplemental Figures**

**Figure S1. *Tg*TR consumption of NAD(P)H with octanal.**

**Figure S2. Kinetics of substrate binding to *Tg*TR.**

**Figure S3. *Tg*TR Tm in the presence of cofactor or substrate.**

**Figure S4. Binding affinity to *Tg*TR by ITC.**

**Figure S5. SDS-PAGE gels of purified protein constructs.**

**Figure S6.** **Analysis of ACP-derivatives by LCMS.**

**Figure S7. *In vitro* investigation of *Tg*KS5 domain.**

**Figure S8. TR sequence alignments.**

**Figure S9. Overview of the *T. gondii* lifecycle**

**Table S1.** Amino acid and nucleotide sequences for all plasmid constructs.

| **Name** | **Sequence** |
| --- | --- |
| pMAL-c2X MBP Tag-Factor Xa-*Tg*TR-6xHis | MKIEEGKLVIWINGDKGYNGLAEVGKKFEKDTGIKVTVEHPDKLEEKFPQVAATGDGPDIIFWAHDRFGGYAQSGLLAEITPDKAFQDKLYPFTWDAVRYNGKLIAYPIAVEALSLIYNKDLLPNPPKTWEEIPALDKELKAKGKSALMFNLQEPYFTWPLIAADGGYAFKYENGKYDIKDVGVDNAGAKAGLTFLVDLIKNKHMNADTDYSIAEAAFNKGETAMTINGPWAWSNIDTSKVNYGVTVLPTFKGQPSKPFVGVLSAGINAASPNKELAKEFLENYLLTDEGLEAVNKDKPLGAVALKSYEEELAKDPRIAATMENAQKGEIMPNIPQMSAFWYAVRTAVINAASGRQTVDEALKDAQTNSSSNNNNNNNNNNLGIEGRISSVFLTGATGFIGSQVLFHLLQVKRQDPKLGTEDKKLTIYCLVRARDRAHGIYRIRDSLVGRGIEWKHEFDQQVIPVIGNLEEGENMGLGPKQMQYLEQTVDAVYHAADYVNFAAPYDALRKSNVLSLIPILKLCTTYVAKPLHLVSNFAHHLQYFAGFSEDLNIAVDETVSPPLSPASIDRLEQQMPAGIIGYPWTKWAVEEIIGRMKDVLHERCEEEGQRVSMEESEKNDILSKFQVTVYRLPNSCVYYNNGYTNFANASLSLVLASAQERMIPPGVLPVGAPFLTTPVDLITGILVKLSMTEGKRHHVYHLVSLRVARRKHVVQAIKWLFPDSVECTADELFRRIDENKENSPAHPLRAAMRFWRRYWYSSDTDRDSPFPVAVNNVEDDLPGVMATFPSLASLEHHHHHH* |
|  | ATGAAAATCGAAGAAGGTAAACTGGTAATCTGGATTAACGGCGATAAAGGCTATAACGGTCTCGCTGAAGTCGGTAAGAAATTCGAGAAAGATACCGGAATTAAAGTCACCGTTGAGCATCCGGATAAACTGGAAGAGAAATTCCCACAGGTTGCGGCAACTGGCGATGGCCCTGACATTATCTTCTGGGCACACGACCGCTTTGGTGGCTACGCTCAATCTGGCCTGTTGGCTGAAATCACCCCGGACAAAGCGTTCCAGGACAAGCTGTATCCGTTTACCTGGGATGCCGTACGTTACAACGGCAAGCTGATTGCTTACCCGATCGCTGTTGAAGCGTTATCGCTGATTTATAACAAAGATCTGCTGCCGAACCCGCCAAAAACCTGGGAAGAGATCCCGGCGCTGGATAAAGAACTGAAAGCGAAAGGTAAGAGCGCGCTGATGTTCAACCTGCAAGAACCGTACTTCACCTGGCCGCTGATTGCTGCTGACGGGGGTTATGCGTTCAAGTATGAAAACGGCAAGTACGACATTAAAGACGTGGGCGTGGATAACGCTGGCGCGAAAGCGGGTCTGACCTTCCTGGTTGACCTGATTAAAAACAAACACATGAATGCAGACACCGATTACTCCATCGCAGAAGCTGCCTTTAATAAAGGCGAAACAGCGATGACCATCAACGGCCCGTGGGCATGGTCCAACATCGACACCAGCAAAGTGAATTATGGTGTAACGGTACTGCCGACCTTCAAGGGTCAACCATCCAAACCGTTCGTTGGCGTGCTGAGCGCAGGTATTAACGCCGCCAGTCCGAACAAAGAGCTGGCAAAAGAGTTCCTCGAAAACTATCTGCTGACTGATGAAGGTCTGGAAGCGGTTAATAAAGACAAACCGCTGGGTGCCGTAGCGCTGAAGTCTTACGAGGAAGAGTTGGCGAAAGATCCACGTATTGCCGCCACTATGGAAAACGCCCAGAAAGGTGAAATCATGCCGAACATCCCGCAGATGTCCGCTTTCTGGTATGCCGTGCGTACTGCGGTGATCAACGCCGCCAGCGGTCGTCAGACTGTCGATGAAGCCCTGAAAGACGCGCAGACTAATTCGAGCTCGAACAACAACAACAATAACAATAACAACAACCTCGGGATCGAGGGAAGGATTTCAAGCGTTTTCTTAACCGGTGCGACTGGTTTTATCGGCTCGCAGGTGCTCTTTCATCTGCTGCAAGTTAAAAGACAAGACCCTAAGCTGGGTACAGAGGACAAGAAGCTGACTATATATTGTCTTGTCAGAGCGCGTGATCGGGCTCATGGAATCTATAGAATCCGGGACTCTTTAGTTGGTCGTGGCATAGAGTGGAAGCATGAGTTTGACCAGCAGGTGATACCCGTAATTGGAAACTTGGAAGAGGGCGAAAACATGGGGCTGGGCCCTAAGCAAATGCAATACTTAGAGCAAACGGTGGACGCCGTATACCACGCAGCAGACTACGTCAATTTCGCTGCACCATACGACGCGCTTCGTAAATCAAACGTGCTGTCTCTTATACCGATCCTTAAGTTGTGTACTACGTATGTGGCCAAGCCTCTTCACTTAGTCTCGAACTTCGCTCACCATCTCCAATACTTTGCAGGTTTCTCCGAGGATCTTAATATTGCGGTTGACGAGACGGTATCTCCACCCTTATCACCTGCATCTATAGACCGCCTGGAACAACAAATGCCAGCTGGTATCATCGGTTATCCGTGGACCAAGTGGGCGGTTGAGGAAATAATAGGTAGAATGAAGGACGTCTTGCACGAAAGATGCGAGGAAGAGGGACAGCGTGTCAGTATGGAAGAGAGCGAGAAGAACGACATTCTGTCCAAATTTCAGGTCACAGTTTACCGTTTACCGAACTCTTGTGTCTATTATAATAACGGGTATACAAATTTTGCTAATGCGTCACTGTCTCTCGTGTTGGCTTCCGCGCAGGAACGAATGATACCGCCAGGCGTACTCCCTGTGGGTGCTCCGTTCCTGACTACCCCCGTTGATCTGATAACGGGCATTCTGGTCAAGCTCAGTATGACCGAGGGTAAGCGCCATCACGTCTACCATTTAGTTAGTCTGCGTGTAGCCCGGCGCAAACACGTAGTACAAGCCATTAAGTGGTTATTTCCAGACTCAGTCGAATGCACGGCCGACGAGCTTTTCAGAAGAATCGACGAGAACAAAGAGAACAGCCCGGCACATCCATTAAGAGCGGCCATGCGGTTTTGGAGACGTTACTGGTATAGTTCCGATACCGACCGAGACAGTCCATTTCCTGTTGCTGTCAACAACGTTGAGGACGACTTGCCGGGTGTCATGGCGACTTTTCCGTCACTTGCCTCCCTCGAGCACCACCACCACCACCACTGA |
| pET21a(+) T7 Tag-*Tg*ACP4-6xHis | MASMTGGQQMGRGSMHGSMDIQGVESAVIDVARRLCGAEGIVSASTEISSLGLDSLGAVEFRNALQEALKIKLPASLLVEYACLKDIIQFIMKLAAALEHHHHHH* |
|  | ATGGCTAGCATGACTGGTGGACAGCAAATGGGTCGCGGATCCGAGCCTGACCAGATGGCGAACGAGGTAGTGAATACTGTAGATGAGGAGGCGGTACAGACCCTAGTTTTACAAACTGTACGTCAAATACTGAAGGAGGCTGTGAGTATCCATAGGGACGTCAGCCTGAAAACACTAGGTATCGACAGCCTAGCTGCGGTGGAATTTCGTGATTCCTTACAGGAGTCATTGGGCATTAGTCTGCCGGCCAGTTTGATTTTCGATTTCCCCACGATCGACGATGTATGTGGCTTTCTGGCCCAGAAACTGAAGCTTGCGGCCGCACTCGAGCACCACCACCACCACCACTGA |
| pET15b 6xHis-TEV- Sfp | MGSSHHHHHHSSGENLYFQGHMKIYGIYMDRPLSQEENERFMTFISPEKREKCRRFYHKEDAHRTLLGDVLVRSVISRQYQLDKSDIRFSTQEYGKPCIPDLPDAHFNISHSGRWVIGAFDSQPIGIDIEKTKPISLEIAKRFFSKTEYSDLLAKDKDEQTDYFYHLWSMKESFIKQEGKGLSLPLDSFSVRLHQDGQVSIELPDSHSPCYIKTYEVDPGYKMAVCAAHPDFPEDITMVSYEELLLEDPAANKARKEAELAAATAEQ* |
|  | ATGGGCAGCAGCCATCATCATCATCATCACAGCAGCGGCGAAAACCTGTATTTTCAGGGCCATATGAAGATTTACGGAATTTATATGGACCGCCCGCTTTCACAGGAAGAAAATGAACGGTTCATGACTTTCATATCACCTGAAAAACGGGAGAAATGCCGGAGATTTTATCATAAAGAAGATGCTCACCGCACCCTGCTGGGAGATGTGCTCGTTCGCTCAGTCATAAGCAGGCAGTATCAGTTGGACAAATCCGATATCCGCTTTAGCACGCAGGAATACGGGAAGCCGTGCATCCCTGATCTTCCCGACGCTCATTTCAACATTTCTCACTCCGGCCGCTGGGTCATTGGTGCGTTTGATTCACAGCCGATCGGCATAGATATCGAAAAAACGAAACCGATCAGCCTTGAGATCGCCAAGCGCTTCTTTTCAAAAACAGAGTACAGCGACCTTTTAGCAAAAGACAAGGACGAGCAGACAGACTATTTTTATCATCTATGGTCAATGAAAGAAAGCTTTATCAAACAGGAAGGCAAAGGCTTATCGCTTCCGCTTGATTCCTTTTCAGTGCGCCTGCATCAGGACGGACAAGTATCCATTGAGCTTCCGGACAGCCATTCCCCATGCTATATCAAAACGTATGAGGTCGATCCCGGCTACAAAATGGCTGTATGCGCCGCACACCCTGATTTCCCCGAGGATATCACAATGGTCTCGTACGAAGAGCTTTTACTCGAGGATCCGGCTGCTAACAAAGCCCGAAAGGAAGCTGAGTTGGCTGCTGCCACCGCTGAGCAATAA |
| pMAL-c2X MBP Tag-Factor Xa-*Tg*KS5-6xHis | MKIEEGKLVIWINGDKGYNGLAEVGKKFEKDTGIKVTVEHPDKLEEKFPQVAATGDGPDIIFWAHDRFGGYAQSGLLAEITPDKAFQDKLYPFTWDAVRYNGKLIAYPIAVEALSLIYNKDLLPNPPKTWEEIPALDKELKAKGKSALMFNLQEPYFTWPLIAADGGYAFKYENGKYDIKDVGVDNAGAKAGLTFLVDLIKNKHMNADTDYSIAEAAFNKGETAMTINGPWAWSNIDTSKVNYGVTVLPTFKGQPSKPFVGVLSAGINAASPNKELAKEFLENYLLTDEGLEAVNKDKPLGAVALKSYEEELAKDPRIAATMENAQKGEIMPNIPQMSAFWYAVRTAVINAASGRQTVDEALKDAQTNSSSNNNNNNNNNNLGIEGRISAVVGMGCRIPKNATTTEEFWDMLLHEIDAVCEIPMDRFNIDAFYDPDPEQDKCYTREAALLDHPDFFDNTFFNLSDQEVLAIDPQQRVLLEVAYEALFEAGFTKEALLGKDFGVFCAAYNNDFQFTNLTQGKATMCVFDKGEGRGRIPSLGEAAGYPEPGGFMCFIPNRISYSFGLTGPSIGIDAACASSLVAFDAAVTKLKRGACSGALVGGVNIILSPTFFLGGCKAHQFSRAGRCKTFDCNADGLVRGEGCGAAFLLPLDAARETGAFVHAVVRGSATCHYGRSSQITSPNTRALTRVLRLSLQDASTAPSLVRYYEAHGTATVLGDVIEMSAVKDVFQTGRNAAAPLHMGTVHNNIGHLDAAAGIVAFLKTVLCLKHRFVPANIHFKSLHPDIHGIDTQLIKYTRESHSIMTDDVATGVRKTIFGANLAYGMGGSVAAIITESSLEHHHHHH* |
|  | ATGAAAATCGAAGAAGGTAAACTGGTAATCTGGATTAACGGCGATAAAGGCTATAACGGTCTCGCTGAAGTCGGTAAGAAATTCGAGAAAGATACCGGAATTAAAGTCACCGTTGAGCATCCGGATAAACTGGAAGAGAAATTCCCACAGGTTGCGGCAACTGGCGATGGCCCTGACATTATCTTCTGGGCACACGACCGCTTTGGTGGCTACGCTCAATCTGGCCTGTTGGCTGAAATCACCCCGGACAAAGCGTTCCAGGACAAGCTGTATCCGTTTACCTGGGATGCCGTACGTTACAACGGCAAGCTGATTGCTTACCCGATCGCTGTTGAAGCGTTATCGCTGATTTATAACAAAGATCTGCTGCCGAACCCGCCAAAAACCTGGGAAGAGATCCCGGCGCTGGATAAAGAACTGAAAGCGAAAGGTAAGAGCGCGCTGATGTTCAACCTGCAAGAACCGTACTTCACCTGGCCGCTGATTGCTGCTGACGGGGGTTATGCGTTCAAGTATGAAAACGGCAAGTACGACATTAAAGACGTGGGCGTGGATAACGCTGGCGCGAAAGCGGGTCTGACCTTCCTGGTTGACCTGATTAAAAACAAACACATGAATGCAGACACCGATTACTCCATCGCAGAAGCTGCCTTTAATAAAGGCGAAACAGCGATGACCATCAACGGCCCGTGGGCATGGTCCAACATCGACACCAGCAAAGTGAATTATGGTGTAACGGTACTGCCGACCTTCAAGGGTCAACCATCCAAACCGTTCGTTGGCGTGCTGAGCGCAGGTATTAACGCCGCCAGTCCGAACAAAGAGCTGGCAAAAGAGTTCCTCGAAAACTATCTGCTGACTGATGAAGGTCTGGAAGCGGTTAATAAAGACAAACCGCTGGGTGCCGTAGCGCTGAAGTCTTACGAGGAAGAGTTGGCGAAAGATCCACGTATTGCCGCCACTATGGAAAACGCCCAGAAAGGTGAAATCATGCCGAACATCCCGCAGATGTCCGCTTTCTGGTATGCCGTGCGTACTGCGGTGATCAACGCCGCCAGCGGTCGTCAGACTGTCGATGAAGCCCTGAAAGACGCGCAGACTAATTCGAGCTCGAACAACAACAACAATAACAATAACAACAACCTCGGGATCGAGGGAAGGATTTCAGCGGTCGTTGGTATGGGGTGCAGGATTCCGAAGAACGCGACGACGACGGAGGAGTTCTGGGATATGCTACTTCACGAGATCGACGCTGTATGCGAGATTCCAATGGATCGTTTCAATATTGATGCATTCTACGACCCCGATCCGGAACAAGACAAGTGCTACACGAGGGAGGCAGCCCTTCTGGACCATCCAGATTTCTTTGACAACACATTCTTCAATCTAAGCGATCAGGAAGTGTTGGCAATCGACCCCCAGCAAAGAGTATTATTAGAGGTAGCATACGAAGCACTGTTTGAAGCGGGTTTCACTAAGGAAGCATTGTTGGGCAAGGACTTCGGCGTATTTTGCGCGGCCTACAACAACGACTTTCAGTTCACAAATCTGACACAGGGAAAAGCCACGATGTGCGTGTTTGATAAAGGGGAGGGGAGGGGACGTATCCCCTCATTGGGAGAAGCGGCCGGTTACCCCGAACCAGGTGGGTTTATGTGTTTCATACCTAACCGTATCTCTTACTCATTTGGGTTGACCGGGCCATCTATAGGCATAGATGCTGCGTGTGCTAGCTCTTTGGTAGCTTTTGACGCCGCTGTGACGAAGTTAAAGCGTGGCGCTTGCAGCGGCGCCCTGGTGGGCGGAGTGAACATTATTTTAAGCCCCACATTTTTTCTAGGCGGCTGCAAAGCCCATCAGTTCTCTAGAGCCGGCCGTTGTAAAACCTTCGACTGCAACGCAGATGGTTTAGTAAGGGGGGAGGGTTGTGGCGCTGCGTTTCTGCTACCGCTGGATGCTGCTAGAGAGACTGGAGCGTTTGTGCATGCTGTAGTACGTGGTTCAGCAACATGTCATTATGGCAGATCCAGTCAGATAACCTCCCCAAACACCAGAGCCTTAACCAGAGTGTTGCGTCTTTCATTGCAGGACGCGAGCACAGCCCCATCCTTGGTCAGGTATTACGAAGCCCACGGAACCGCAACGGTACTAGGAGATGTTATAGAGATGTCCGCAGTCAAGGACGTATTTCAGACGGGGAGGAACGCAGCCGCCCCTCTACATATGGGCACAGTTCACAATAATATAGGACATCTAGATGCGGCGGCTGGCATTGTGGCGTTCTTAAAAACGGTTCTGTGCTTGAAACATCGTTTCGTTCCGGCCAACATTCACTTTAAGTCTTTGCATCCAGATATTCACGGTATAGACACCCAGCTTATTAAGTATACCCGTGAATCACACTCTATTATGACAGATGATGTCGCTACTGGGGTGCGTAAAACGATCTTCGGGGCCAATTTGGCCTACGGAATGGGAGGAAGCGTCGCCGCGATAATAACTGAGAGCTCCCTCGAGCACCACCACCACCACCACTGA |

**Table S2.** Domain name and amino acid sequence of TE/TR domains used for phylogenetic trees.

| **Name** | **Amino Acid Sequence** |
| --- | --- |
| Cinnamomum camphora TE (FAS) | MATTSLASAFCSMKAVMLARDGRGMKPRSSDLQLRAGNAQTSLKMINGTKFSYTESLKKLPDWSMLFAVITTIFSAAEKQWTNLEWKPKPNPPQLLDDHFGPHGLVFRRTFAIRSYEVGPDRSTSIVAVMNHLQEAALNHAKSVGILGDGFGTTLEMSKRDLIWVVKRTHVAVERYPAWGDTVEVECWVGASGNNGRRHDFLVRDCKTGEILTRCTSLSVMMNTRTRRLSKIPEEVRGEIGPAFIDNVAVKDEEIKKPQKLNDSTADYIQGGLTPRWNDLDINQHVNNIKYVDWILETVPDSIFESHHISSFTIEYRRECTMDSVLQSLTTVSGGSSEAGLVCEHLLQLEGGSEVLRAKTEWRPKLTDSFRGISVIPAESSV |
| Carthamus tinctorius TE (FAS) | MLSRPLPTTAAAATTTTNNCNGVNSRGALPHSRSVGFASIRKRSTGSLCNSPPRTVAPVMAVRTGEQPTGVAVGLKEAEAEVEKSLADRLRMGSLTEDGLSYKERFIIRCYEVGINKTATVETIANLLQEVGGNHAQSVGFSTDGFATTTTMRKLHLIWVTSRMHIEIYRYPAWSDVVEIETWCQSEGRIGTRRDWIMKDHASGEVIGRATSKWVMMNEDTRRLQKVNDDVRDEYLVFCPKTPRLAFPEKNTSSLKKIAKLEDPAEYSTLGLVPRRADLDMNKHVNNVTYIGWVLESIPQEVIDTHELQTITLDYRRECQHDDIVDSLTSSESLLDDAAISKLEGTNGSSVPKKDETDLSRFLHLLRSSGDGLELNRGRTEWRKKPAKK |
| Brassicajuncea TE (FAS) | MLKLSCNVTNHLHTFFFSSDSSLFIPGNRRTIAVSSSQPRKPALDPLRAVISADQGSISPVNSYTPADRFRAGRLMEDGYSYKEKFIVRSYEVGINKTATVETIANLLQKVACTHVQNVGFSTDGFATTLTMRKLHLIWVTARMHIEIYRYPAWSDVVEIETWCQSEGRIGTRRDWILRDSATNEVIGRATSKWVMMNQDTRRLQRVTDEVRDEYLVFCPREPRLAFPEENNSSLKKIPKLEDPAQYSMLELKPRQADLGMNQHVNNVTYIGWVLESIPQEIIDTHELQVITLDYRRECQQDDIVDSLTTSEIPDDPISKLTGTNGSATSRVQGHNESQFLHMRLSENGQEINRGRTQWRKKSPR |
| Arabidopsis thaliana TE (FAS) | MLKLSCNVTDSKLQRSLLFFSHSYRSDPVNFIRRRIVSCSQTKKTGLVPLRAVVSADQGSVVQGLATLADQLRLGSLTEDGLSYKEKFVVRSYEVGSNKTATVETIANLLQEVGCNHAQSVGFSTDGFATTTTMRKLHLIWVTARMHIEIYKYPAWGDVVEIETWCQSEGRIGTRRDWILKDSVTGEVTGRATSKWVMMNQDTRRLQKVSDDVRDEYLVFCPQEPRLAFPEENNRSLKKIPKLEDPAQYSMIGLKPRRADLDMNQHVNNVTYIGWVLESIPQEIVDTHELQVITLDYRRECQQDDVVDSLTTTTSEIGGTNGSATSGTQGHNDSQFLHLLRLSGDGQEINRGTTLWRKKPSS |
| Equisetin Fusarium heterosporum TR (PKS-NRPS) | VLLTGSSGYLGRHLLSSLLNDHRVAQVHCLCRNLNDHQVVENPSSKVRVLQSDLAQHKLGLPDSTYSQLATEVDVIIHCAANRSFWDRYEALKADNLESTKELVKLVVSSGRAIPLHFLSSGAVIKYNSGLAPPADGGDGYVATKWASEAFLRQAVDSINLPVFSHRPVACESVQQSEEEDISILNELIQIVKLLGCRPSFDGVGGFVDVMPVNEVVEAI |
| Hancockiamides Aspergillus hancockii TR (NRPS) | RVFLTGATGFVGAFLLSDMLKMPGIHQVGCLVRAPDEATGVRRLRHALEKYNLWREEYLPKLLPLCGKLEDPWLGLGEQRFREIADWASVIFHLGALVNYTQPYSWHRPANIEGTVNVVRLACTGRSKALHYCSSISCFGPTGIINGTKVVHEDGALMPHLNALPYDHGYAQSQWVAEELLRRLIHRRFPIAVYRPGFITGHSETGACNPDDFFSRLIRACSSIGCYPGLPNQRKEFVPIDYVTSTMIHIASSSLSLGHAFHIVPPTREESPEMNDTMSLIGELTGTSIQPVSYREWIEQLSSTKDLSLQPLLPMLAEVVIDGMTRWEMYENMPTYENTNTLRALASCPDLPKFPMVDEALLRKYLDYLA |
| Nostocaceae sp. TR (NRPS) | KVFLTGGTGFLGAFLIRELLQQTQADVYCLVRAADAQAGKAKIQTNLEGYAIWQEEYESRIIPVVGDLAEPLLGLSSTQFQALAAEIDTIYHSGALLNYVFPYSALKAANVLGTQEVLRLACQIKVKPVHYVSSVAVFESPVYAGKVVKESDDFSHWEGIFLGYSQTKWVAEKLVKIAGDRGLPVTIHRPPLIGGDSETGIGNTHDFINLMVKGCLQMGCFPDVEYMLDMSPVDYVSKAIVHLSMQKESLGKAFHLQHPEPVSLSVLVDWVRSFGYEIKTLPYEEWQAELINNATSVDNPLYTLRPFLLERWSDEQLTIPDLYLKAKRPHISCQDTLKALAGTSIGCPPIDGKLFMTYTAYLIQTGFL |
| Azinomycin Streptomyces sahachiroi TR (PKS-NRPS) | VLLTGATGFVGRFVLAELLAAGARVICLLRGGTARREELVAGMADLGLWHEEHAARLELVDGDIAEPGLGLAGPDRDRLADRAGRIIHAAAWVNHVYPYERLAAANTHCMAGLLELAARGRRSALTVVSTSSVADSAAYPPGSTVPPGPLKALPSAANGYVRSKAVAEQYLHLAAELDVPAAVIRIPSVFGDQRRYQINPADAVWSWCRAMIETSGFPESFAQPGNELFQALPADAVARAV |
| Lyngbyatoxin Lyngbya majuscula TR (NRPS) | MTNPFADPSRDYWVLCNAEGQYSLWPTSLEIPEGWQTAFGPESWQNCLDNVEKNWTDMRPLSLLGQRKCPVPYPFREYVALNMDPIYASLQQRDEMLRISVPYGDDAWLASSYKHVKCVIQDTRFSRETTKYDEARLTPIPIRTSVLGMDPPDHTRLREALAAALTPSHVEQMKPWITSMTEQLIDDFVAQGPPADLVEQFALPFTGHITCELMGIPLEDRPQFKAWCDGFSSTSSLTKEEVEVRMQAMYTYITELVGRRRESPSDDIISKLLQPEDKRLELSEIELIDLATILLLAGYDSTAMELANAIFVLLTHPEQTQLIREQPKLMPQAVQELLRYIPIDAHVTFARYATEDVQVGNTLVRTGDAILASFPSANHDPAIFEDPHTFNIMRPRKPNFGLGTGIHSCAGKLLAVVELEIALSVLIRRLPTLRLAIPAEEVPWQPGSLLRSTSKLPAEW |
| Myxalamid Stigmatella aurantiaca TR (PKS-NRPS) | DVTVEMEADAVLDAEIALGKALPPVTGALRTILLTGATGFLGAFLLEELCRRTDARIYCLVRSKTEQEGMNRIRKNLESYSLWNEALAPRIVPVRGDIGQPLLGLSEKEFQRLSEEIDAIYHNGALVNFLYPYESMRAANVLGTREILRLATRTRIKPLHYVSTVSVLPLGRKAPIREDEPLEGPSSLVGGYAQSKWVAEKLVREASRRGLPVTILRPGRVTGHSRTGAWNTDDLVCRTLKGCVRMGVAPSVDALLDLTPVDYVSSAIVDLSMRPESIGQTYHLVNPQFVRADEMWNYMRAFGYGLRVLPYDQWLSELGSAASSDSELGDLLMFLQQVPPEDRSVGGPRMVVCDSGDTLKALGGTGTSCPSVDASLISTYLSSLVHRGFLKAPE |
| Myxochelin A Stigmatella aurantiaca TR (NRPS) | EQHSVEGEGISKAMLADAELPEEIVPRLPTPGAEAPLAPSPGPAAPLRQVLLTGATGFVGAHLLDQLLRQTQAKVVCLVRARDEAHAMERLREAMTSQRLSTASLSERVLALPADLGQPWLGLSSARFHGLAAECDMILHNAAVVSVVREYGSLQATNVRGTRELLRLAASVRPKPLHYVSTLAVAPQANLSPEVPEAFVPAHPGLRDGYQQSKWAAERLVEQASERGLPVTVYRLGRVSGALDSGIVNPQDLVWRILLAGIPAGALPQLDVGEVWTPVDYVARALVRLSLVPRPGTVFNLTPAPEVRLSEVFGWVQDYGYPVALCPVPEWRTRVAQSTGSAENSTTLAFFDLRAGAAEPTFGLGTIRSERVLQALSDTGISCPRTDRPLLHRYLDYCVGQ |
| Glarea lozoyensis TE (NRPS-PKS) | HLWMVPDGSGSATSYTEISEIGPNVAVWGLNSPYMKVPEEYKCGVIGMASSFISEIKRRQPVGPYLLAGWSAGGVIAFEAVNQLIKNGDEVEQLIIIDAPCPDIIEPLPASLHRWFGEIGLLGDGDLSKLPKWLLPHFAASVTALSNYSAELIDPKKSPFVTAIWCEDGVCHLPTDPRPDPFPYGHAQFLLDNRTDFGPNLWDKYLNGERFVTKHMPGNHFSMMK |
| Pochonia chlamydosporia TE (PKS) | LIADGTGSIATYLHLPPHINTKMTIYGVDSPYLHCPSRLTPDVGIPGIAKLIVDELVKRQPQGVPFWLGGFSGGAMIAYEIARQLSALGHVIDSLLLIDMCPPRQIQAQRYDDELGLAMFDAISGNDDSGVWESSDKTHQHLKA |
| Naphthopyrone Aspergillus nidulans TE (PKS) | TLFLFPDGSGSATSYATIPGVSPNVAVYGLNCPYMKAPEKLTCSLDSLTTPYLAEIRRRQPTGPYNLGGWSAGGICAYDAARKLVLQQGEIVETLLLLDTPFPIGLEKLPPRLYSFFNSIGLFGEGKAAPPAWLLPHFLAFIDSLDAYKAVPLPFNEQEWKGKLPKTYLVWAKDGVCPKPGDPWPEPAEDGSKDPREMVWLLSNRTDLGPNGWDTLVGKENIGGITVIHDANHFTMTKGEKAKELATFMKN |
| Sterigmatocystin Aspergillus nidulans TE (PKS) | ILFMLPDGGGSASSYLTIPRLHADVAIVGLNCPYARDPENMNCTHQSMIQSFCNEIKRRQPEGPYHLGGWSSGGAFAYVTAEALINAGNEVHSLIIIDAPVPQVMEKLPTSFYEYCNNLGLFSNQPGGTTDGTAQPPPYLIPHFQATVDVMLDYRVAPLKTNRMPKVGIIWASETVMDEDNAPKMKGMHFMVQKRWDFGPDGWDVVCPGAVF |
| Gulmirecin A Pyxidicoccus fallax TE (PKS) | RLLCFPHGGAGASAFQGWWQQFPDTVEVLTFQPPGREERLQEEPERDMGAYASAIVEELQPLLDLPFAVFGYSLGGLTGFATVAELRRRNRVTPRHLFVSSAFAPHIPFSETTLARVSTQTSALKAMAFYQSTPDSVFKDPEMQALLARSMDADNSVVESYRFGAEKPLDCPITAFAGRQDDLAPLHEQLRWKELTTQAFELRPFEGDHFFFLRDK |
| Aflaxtoxin Aspergillus ochraceoroseus TE (PKS) | RTLFMLPDGGGSASSYITIPRLQSDVAIVGLNCPYARDPENMQCTHQAMIQSFCNEIKRRQPQGPYHLGGWSSGGAFAYVTAEALVNMGEDVHSLIIIDAPVPQVMEKLPTSFYEYSNNIGLFANQPGSNTDGAAQPPPYLIPHFQATVDVMLDYRVAPLKTPRMPKVGIIWAATTVMEEANAPKMKGMHFMVQKRTDFGPDGWDTVLPGAEF |
| Aureothin Streptomyces thioluteus TE (PKS) | KLICFPTFAGGSGAHQYARLAAVSGGERDIWVLPAPGFTWQESLPASVETLARLQADSVEACADGGPVVLLGYSAGGWIAQATATALQERGIAPAAVVLLDSHRPDSAMLPHLHAHIDGAREDVLWTAGTGDDAYLTAMAHYAHLFHTWTPKETGTPTLLVRASDRPFDEAPTDWQPHWPLPHTAVDTAGTHFSIIREHSGPTLQAVHAWL |
| Pseudomonas cannabina TE (PKS) | TYQRLSRELKGHYDMATLPLPGYDNGPLPASAEAVAVALAHAVEACAAGRPFTLLGFSTGGLVAYAVAELLQQRGIHPAALCLIDTYPPAAMRSAVLKEVLSDWLESRSDFWSTDDDGLSAMAYYLELFGRWSPLPLEAPVLLLQAEQASAAGASDTWPQLWPRLTQVVKTSGRHFALLSDEVQQTAQHLI |
| Streptomyces aizunensis TE (PKS) | LVCFSSILSISGPHQYARFASAFRGRRDVHALGAPGFLRGEQLPSATDAVIEAQAEAVLRHADGAPFVLLGHSSGGMLAHAVAGRLESEGVFPQALVMIDIYSHDDDAIIGIQPGLSEGMDERQDTYVPVDDNRLLAMGAYFRLFGGWKPEVVKTPTLLVRAGERFFDWTRSTDGDWRSYWDLDHTALDVPGNHFTMMEEHAPTTAQAVEGWL |
| Erythromycin A EryAIII TE (PKS) | VCCAGTAAPSGPHEFARLAAALDGSWSVLGLPQPGYLAGEPLPASMEALAAAQAATVLRAVAARPVVLVGHSAGGLMSHALATALTQQGHRPDGVVLLDSYPPGRQDAVEGWIDVLTRRLLDLETVPLDDTRLTAMGAYDRMLGGWVPDDQQVATLLVRAVDPIAAWPDDDWRASWPFAHATADVPGDHFTMMQEHAGQVAARITDW |
| Vitrella brassicaformis TR (Putative PKS) | VRHVLLTGVTGFVGRLQLVCLLNRPGGDNLTVHCLVRANNEKHAMQRIREACIEAKVWKEEYASRIVALLGDFTKEDLGVGEDKFMELCRTIDMVHHTGGDVGLLSNYARLRATNTLALKGVLRLCTTYRLKSLHHASTLGIFPAFFACFSQDFDDRLITEDSSPDPKEMQKFFPPLRMGYPWSKWAAEQVLRKARDMGLPVCVYRLPNTFVAWRTGYTNKGDYATALTISSIEEGMFPVGASAAPFTPVDTIAEMLVELSFLERRKHWLYHLINTTVVTQAQSKAWADELGIKYYGVKIDTFLDAVKKHGPESPVFKFVPLMQHWRSYWFDCDERTEPLPIRTQNIFDELP |
| Toxoplasma gondii TgPKS2 TgTR (PKS) | SVFLTGATGFIGSQVLFHLLQVKRQDPKLGTEDKKLTIYCLVRARDRAHGIYRIRDSLVGRGIEWKHEFDQQVIPVIGNLEEGENMGLGPKQMQYLEQTVDAVYHAADYVNFAAPYDALRKSNVLSLIPILKLCTTYVAKPLHLVSNFAHHLQYFAGFSEDLNIAVDETVSPPLSPASIDRLEQQMPAGIIGYPWTKWAVEEIIGRMKDVLHERCEEEGQRVSMEESEKNDILSKFQVTVYRLPNSCVYYNNGYTNFANASLSLVLASAQERMIPPGVLPVGAPFLTTPVDLITGILVKLSMTEGKRHHVYHLVSLRVARRKHVVQAIKWLFPDSVECTADELFRRIDENKENSPAHPLRAAMRFWRRYWYSSDTDRDSPFPVAVNNVEDDLPGVMATFPSLA |
| Eimeria acervulina TR (Putative PKS) | VSEIFLTGATGLTGSRILLSLLEQQVDGRNDGRKGFPRIYCLVQARDRMHAMYRIVDAVVGRGNQWKQRFFEQVVPVVGDLRAQRLGLSQKTFDELANKVEAVYHAGRVVDFALPYEAIRKANVLSLLPLVELCTTGKAKHLHVLSDFAAHIQYFGAFAGDLNQPILEELSVPQPLMDRMENQMPASVMGYPWARWAVEEVLAKCQSWMGNLHASAGGDFHDIKTKFRFTIYRLPNSAVYFDNGRVDFLNPFFTITMAALQQGLIPPGVLPVGPPFLSTPIDISADILVKLSRLPHRPTVMHIVNPVGVRREAVNQTLLALKIPFRESCEDEFLRGIENNQNGSAAYTLLPVMRFWRRYWFSEDLDRSDAFPLSTANVEACLPNALNQFP |
| Eimeria catenella TR (Putative PKS) | VSEIFLTGATGLTGSRILFGLLEQRIDCGKEVQSRFPRIYCLVQARDRMHAMYRIVDAAVGRGQKWKARFFEQIVPVVGDLKAPRMGLSSKTFETLATRVEAVYHAARVVDFAQPYEKIRKANVLSLLPLIELCTTGIAKHFHVLSDFAAHIQYFGAFAGDLNVSLPEQLSVPDELVCRMENQMPASIIGYPWARWAVEEVLLKCQQRLDELKKSATGSSHDIENKFAFTVYRIPNSAVYFNNGRIEFSNSFFAIVMAALQEGIVPPGVLPVGPPFLTTPIDISADILVNISRCPERPTVIHIVNPTGVRRSAIIEALQSLGIPFRESCEDELFRGIEKNQNGSCAYSLLPIMRFWRRYWFSEDLDREAFPIEIANVEACLPHALKRFPS |
| Eimeria necatrix TR (Putative PKS) | SEIFLTGATGLTGSRILFSLLEQRIDCGKGAQSRFPRIYCLVQARDRMHAMYRIIDAAVGRGQKWKARFFEQIVPVVGDLKAPRMGLSSKTFETLATRVEAVYHAARVVDFAQPYEKIRKANVLSLLPLIELCTTGIAKHFHVLSDFAAHIQYFGAFAGDLNVSLPEKLSVPDDLVGRMENQMPASIIGYPWARWAVEEVLSKCQQRLDELKKSATGSSHDIENKFAFTVYRIPNSAVYFNNGRIEFSNSFFAISMAALQEGIVPPGVLPVGPPFLTTPIDISADILVDISRCPERPTVIHIVNPTGVRRSAIIQALQSFGIPFRESCEDELLRGIEKNQNGSCAYSLLPVMRFWRRYWFSEDLDREAFPIETANLEACLPNALKRFPS |
| Eimeria mitis TR (Putative PKS) | VSEIFLTGATGLTGSQILLSLLEQQIDCGRDGQKRFPRIYCLVQARDRMHAMYRIVDAVIGRGGQWKQRYFEQVVPVVGDLRAPRLGLSQKTFDALTSTVEAVYHAGRVVDFALPYEAIRKANVLSLLPLVEMCTTGKAKHLHVLSDFAAHIQYFAAFAGDLNQPISEELSVSQPLMDRMENQMPASIMGYPWARWAVEEVLAKCQDWIDKLHASAADDLQDAKTKFNFTIYRLPNSAVYFNNGRVEFLNPFFAITMTALQQGVVPPGVLPVGPPFLTTPIDISADILVRISRRLHRPTVIHIVNPVGVRREALNQTLLALKIPFRESCEDEFLRGIEKNQNSSAAYTLLPVMRFWRLYWFSEDLDRKDAFPIVTANIEACLPNALKQFP |
| Cryptosporidium ryanae TR (Putative PKS) | LLTGVTGFVGRILLIKLAEQFKDMKIVCLVRAKDEETALQRIINVCKEAEVWNQSLISRIVVECGDFEKEYLGLSKERYEKLSVEVDVVYHIGGDVNLLSNYKRLRRTNTLSLIGIIDFCCKGRLKHLHFSSTLGQFPAFFSMFTREFENYVVKESECPSTREMSRLFPPTRQGYPWSKWAAEQILEIARDQGLCVSIYRLPNTYIASDTGYTNKTDYATALLISSITEKMFPIGSSTAPLTPVNTICDIIIAASKKKMRKHWRYNLIDTRVINTNDFEAWGSQLGIKEYKGVLIDEFFTVIKNRGPESPIFKFVPLMKYWRHYWFDNVQRTTPFPVDTSNVLEDFPEI |
| Cryptosporidium bovis TR (Putative PKS) | LLTGVTGFVGRILLIKLAEQFKDMKIVCLVRAKNEEVALQRIINVCEEAEVWDQSLVSRIVVECGDFEKEYLGLSKKRYEELSLEVDVVYHIGGDVNLLSNYKRLRKTNTLSLIGIIDFCCRGRLKHLHFSSTLGQFPAFFSMFTREFENYVVTETECPSTKEMSRLFPPTRQGYPWSKWAAEQILEIARDQGLRVSIYRLPNTYIASNTGYTNKTDYATALLISSITEKMFPIGSSTAPLTPVNTICDIIIAASKKKMRKHWRYNLIDTRVINTSDFELWGSQLGIKEYKGVLIDEFFTVIKNRGPESPIFKFVPLMKYWRHYWFDNVQRTTPFPVDTSHVLEDFPEI |
| Cryptosporidium felis TR (Putative PKS) | KTALLTGVTGFVGRVLLSKLINQFDDLQIVCLVRAKTQEKGMERIISVCEEAEIWDSSFASRIVVECGDFEEEYLGLSKERYYELCAEIDVVYHIGGDVNLLSNYKRLRKTNTLSLTGIIEFCSTTKLKHLHFSSTLGQFPAFFAIFTREFENSVVKETEGPSTREMSRLFPPTRQGYPWSKWAAEQILEAAFKQGLPVSIYRLPNTYVASDTGYTNKTDYATALLIASILEGIFPMGSSTAPLTPVNTICDIIVSASKKKKRLHWRYNLIDTRIVSTKNIEVWASQLGIGNYKGVGVDDFFTAIKERGPESPIFKFVPLMQYWRHYWFDNIERTEPFPVDTSNVSDDIPEI |
| Neospora caninum TR (Putative PKS) | VFLTGATGFIGSQVLFHLLQVKRQDTKLGTDGAKLTIFCLVRARDRAHGMYRIRDSLAGRGIEWNPDFDQQVIPVVGNLEEGENMGLGPKQMQYLEQTVDAVYHAADYVNFALPYDALRKANVLSLVPILKLCTTHVAKPLHLVSNFAHHLQYFAGFSEDLNVAVDETVSPPLTPGFVDRLEQQMPAGIMGYPWTKWAAEEIIG |
| Besnoitia besnoiti TR (Putative PKS) | MRSVFVTGATGFIGSQLLFHLLQARCGDLELGEERGKLQIYCLVRARDRAHALYRIKDAFVGRGIEWDPDYNRRVIPVVGDLEENENMGLGVRQLDFLTRVIDAVYHAADNVNFAAPYDELRKTNVTSLIGILKLCTTFVAKPLHLISNFSHHLQYFAAFSNDLNIPVEETISPPLKPDMVKRLEQQMPAGIVGYPWTKWAVEEIVGRMKDVLQHRCKEDSVDKPDTDAILSKFQVVVYRLPNSCVYYRNGYTNFGNASLALVLACAQERMLPPGVPPVGAPFLTTPVDLTTEIVVKLSMTKERQHDVYHLVSLRAARRDNILQVAKWLFSDISECTVDELLRRVDANKENSPAHNLRAAMRFWRSYWYCPDTDRENAFPIMTGHIEDDIPGITTAFPHLA |
| Cyclospora cayetanensis TR (Putative PKS) | IFLTGATGLTGSQILCSLLEQRTCNRTQKENGFLRIYCLVQARDRMHAMYRIVDAVVGRGQRWKPRYFKQIIPVVGDLKAPRLGLSPQNFATLARTIDAVYHAGRDVDFGLPYEAIRKANVLSLLPLVELCTKGKAKPLHVLSDFAAHIQYFAAFGDDLNRPLPEELSVPRELIDRMENQMPASVVGYPWARWAVEEVLANC |

**Table S3.** *Tg*TR site-directed mutagenesis primers.

| **Primer Name** | **Sequence** |
| --- | --- |
| *Tg*TR N147T FOR | 5´-CTTAGTCTCGACATTCGCTCACCATCTCCAATAC-3´ |
| *Tg*TR N147T REV | 5´-TGAAGAGGCTTGGCCACA-3´ |

**Table S4.** qRT-PCR primers.

| **Primer Name** | **Sequence** |
| --- | --- |
| qPCR *TgGAPDH* FOR | 5´-CAGTCGACGAGCATGCAA-3´ |
| qPCR *TgGADPH* REV | 5´-GCAGACTTCCACAGTACC-3´ |
| qPCR *TgSAG2D* FOR | 5´-TTCCGTGTCTTTCGTCTATCC-3´ |
| qPCR *TgSAG2D* REV | 5´-CGTTAGTTTTGCTTGTGATTCGC-3´ |
| qPCR *TgPKS1* FOR | 5´-AGGAGTTCTTGGGAAGATCC-3´ |
| qPCR *TgPKS1* REV | 5´-CCTGCTTCTGGTTACCTTTG-3´ |
| qPCR *TgPKS2* FOR | 5´-ATCAGGGTGCATAGTCATCG-3´ |
| qPCR *TgPKS2* REV | 5´-CCCGTTAGGTGATGTGATGC-3´ |

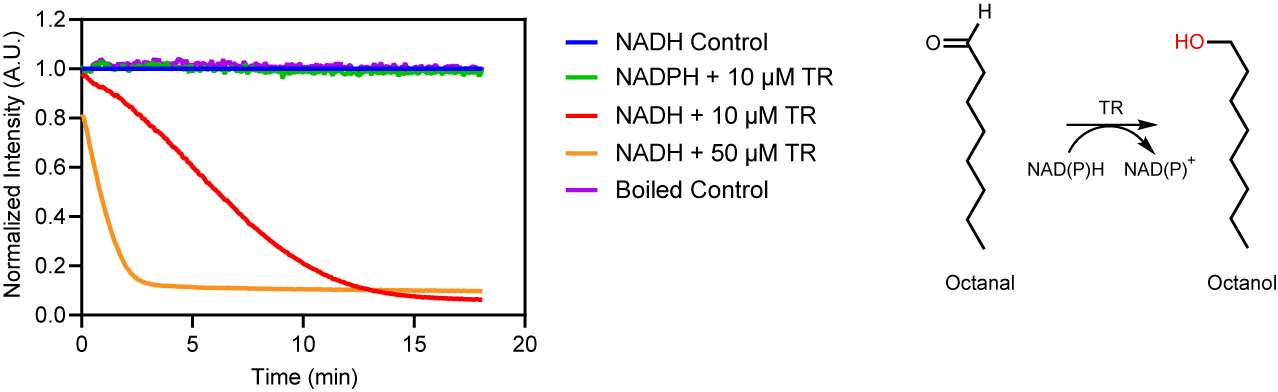

**Figure S1.** *Tg*TR consumption of NAD(P)H with octanal. Normalized fluorescence of NADH control (blue trace), NADPH with 10 µM *Tg*TR (green trace), NADH with 10 µM *Tg*TR (red trace), NADH with 50 µM *Tg*TR (orange trace), and a boiled control containing 10 µM *Tg*TR and NADH (purple trace). All reactions contained 500 µM octanal and 100 µM NAD(P)H. Reaction scheme shown in right panel. All samples were normalized to NADH control.

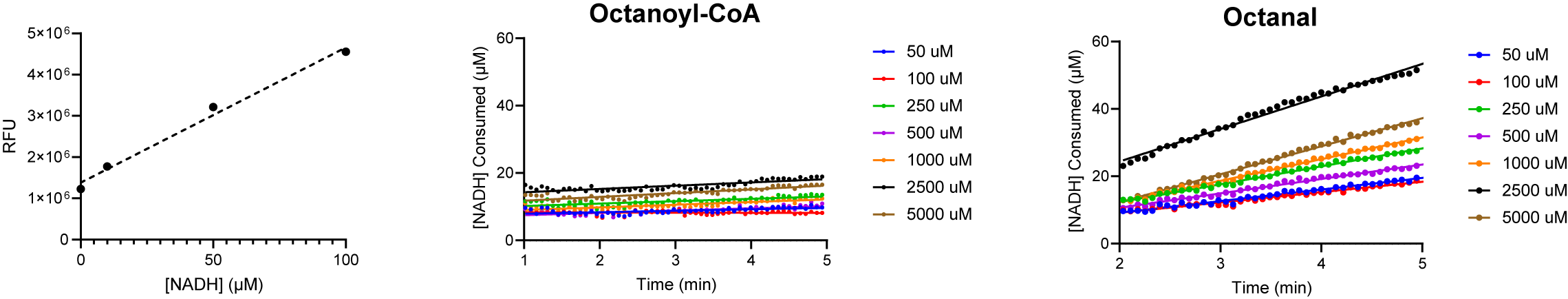

**A**

**B**

**C**

**Figure S2.** Kinetics of substrate binding to *Tg*TR. (A) Calibration curve using NADH standards (0-100 µM), data are shown as the average ± SEM (n=3). (B) Representative data of rate of NADH consumption for varying concentrations of octanoyl-CoA, data are fit to a linear regression to obtain *k*obs. Each reaction contained 10 µM *Tg*TR and 100 µM NADH. (C) Representative data of rate of NADH consumption for varying concentrations of octanal, data are fit to a linear regression to obtain *k*obs. Each reaction contained 10 µM *Tg*TR and 100 µM NADH.

**A**

**B**

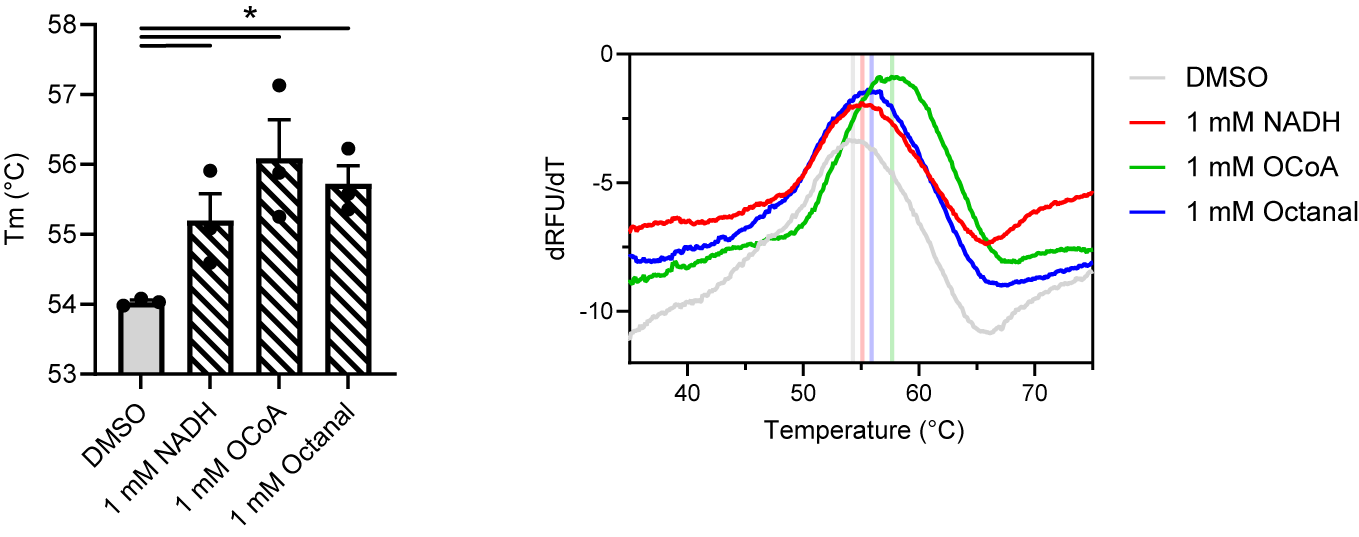

**Figure S3.** *Tg*TR Tm in the presence of cofactor or substrate. (A) Melting temperature (Tm) of *Tg*TR with cofactor and substrate mimics. Tm was calculated by taking the first derivative of the fluorescence values from a thermal shift assay with respect to temperature. Data are shown as the average ± SEM (n=3). * p < 0.05. (B) Representative derivative data used to calculate the Tm for each condition. Tms are indicated by vertical lines.

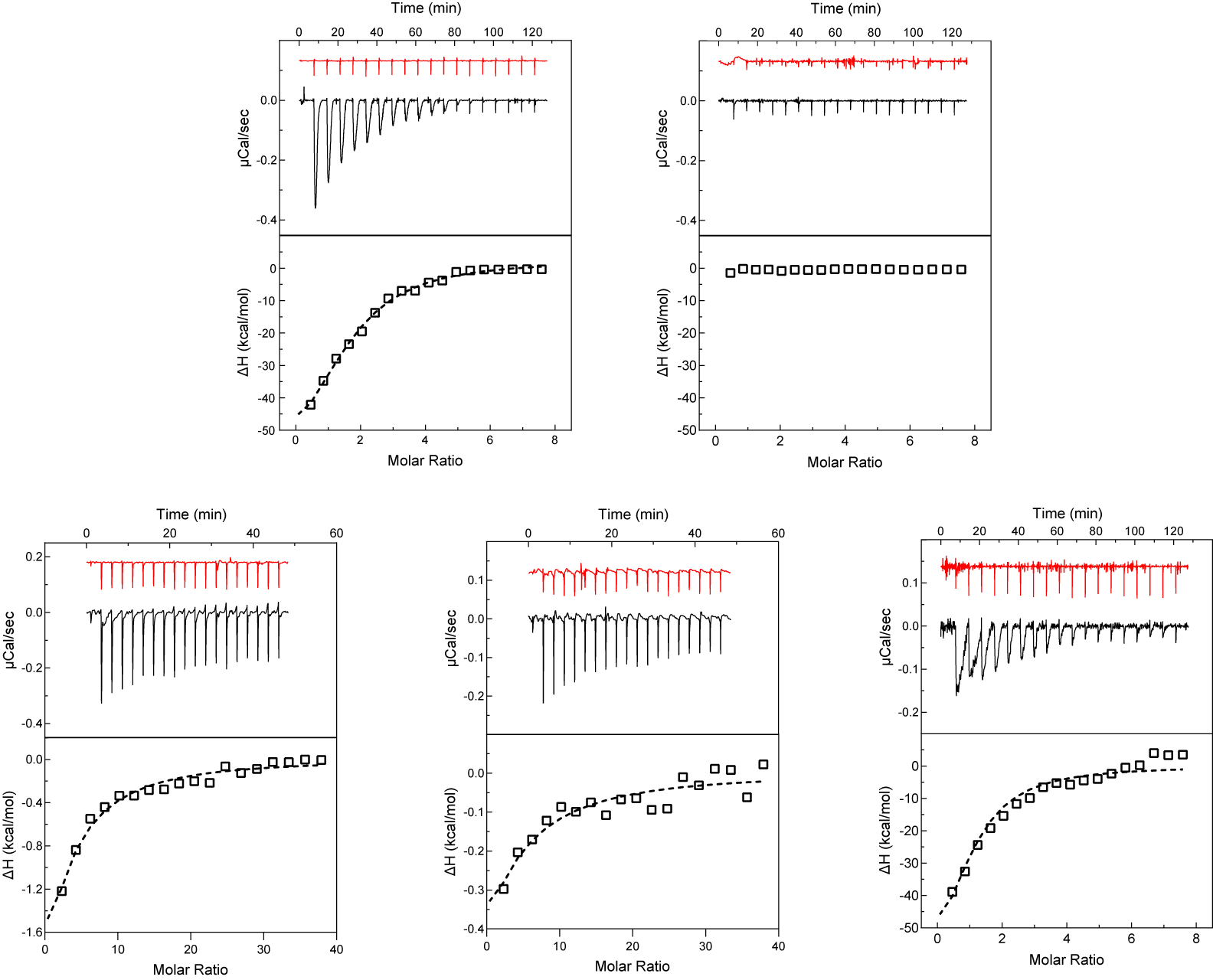

**D**

**E**

**C**

**A**

**B**

**Figure S4.** Binding affinity to *Tg*TR by ITC. Representative ITC thermograms (top panels) and isotherm plots (bottom panels) of injections of (A) 400 µM NADH into *Tg*TR, (B) 400 µM NADPH into *Tg*TR, (C) 2000 µM octanoyl-CoA into *Tg*TR with excess NADH, (D) 2000 µM octanal into *Tg*TR with excess NADH, and (E) 400 µM holo-*Tg*ACP4 into *Tg*TR with excess NADH. Control dilutions into buffer are show in each upper panel (red).

**D**

**C**

**B**

**A**

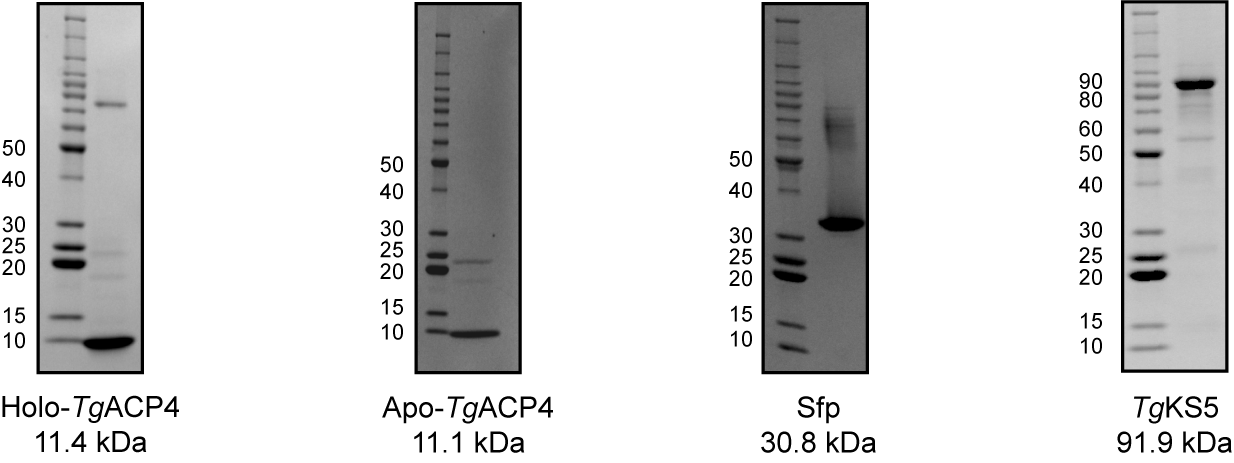

**Figure S5.** SDS-PAGE gels of purified protein constructs. (A) SDS-PAGE gel of holo-*Tg*ACP4, protein ladder is shown in first lane. (B) SDS-PAGE gel of apo-*Tg*ACP4, protein ladder is shown in first lane. (C) SDS-PAGE gel of Sfp, protein ladder is shown in first lane. (D) SDS-PAGE gel of *Tg*KS5, protein ladder is shown in first lane.

**B**

**A**

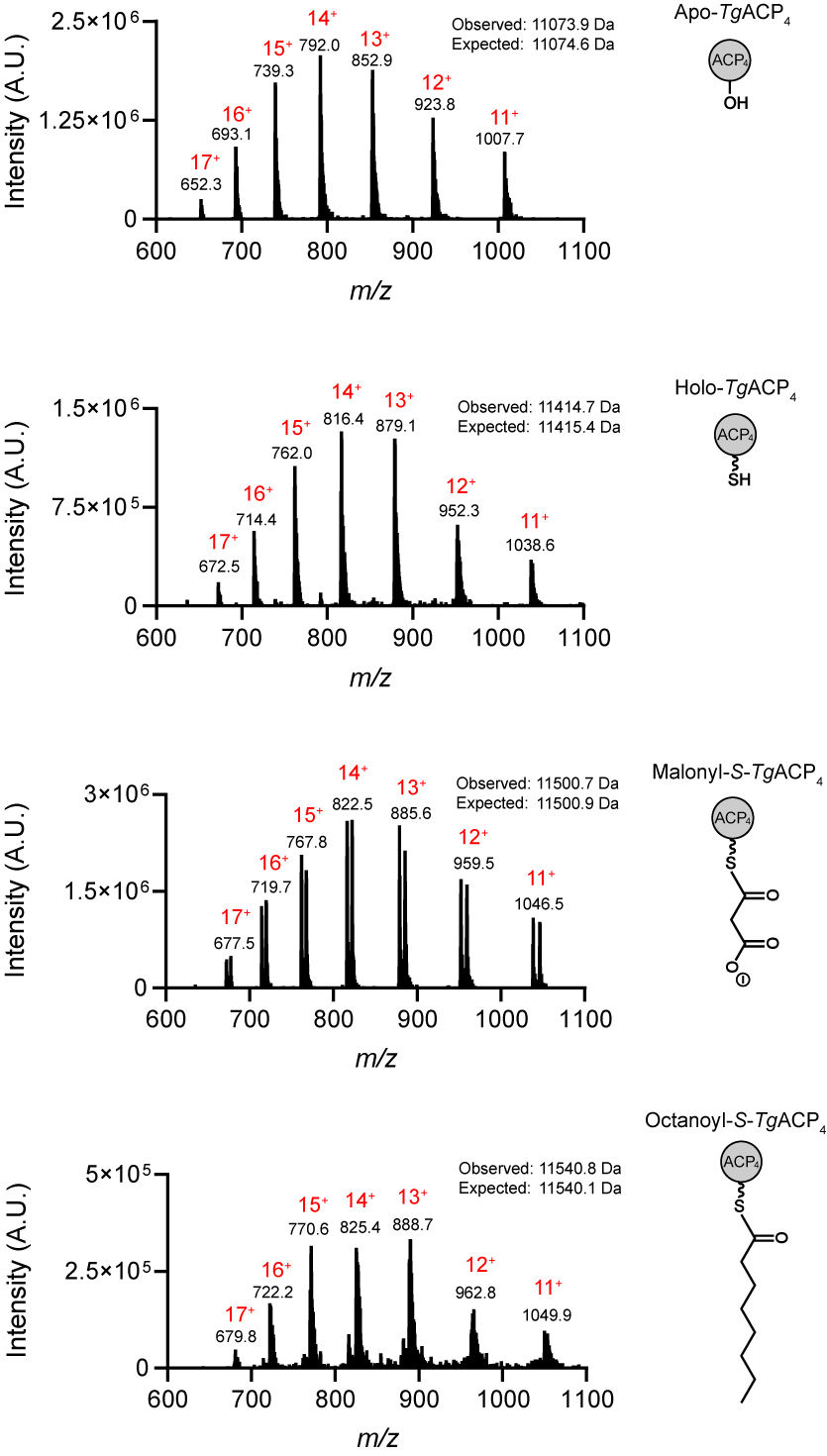

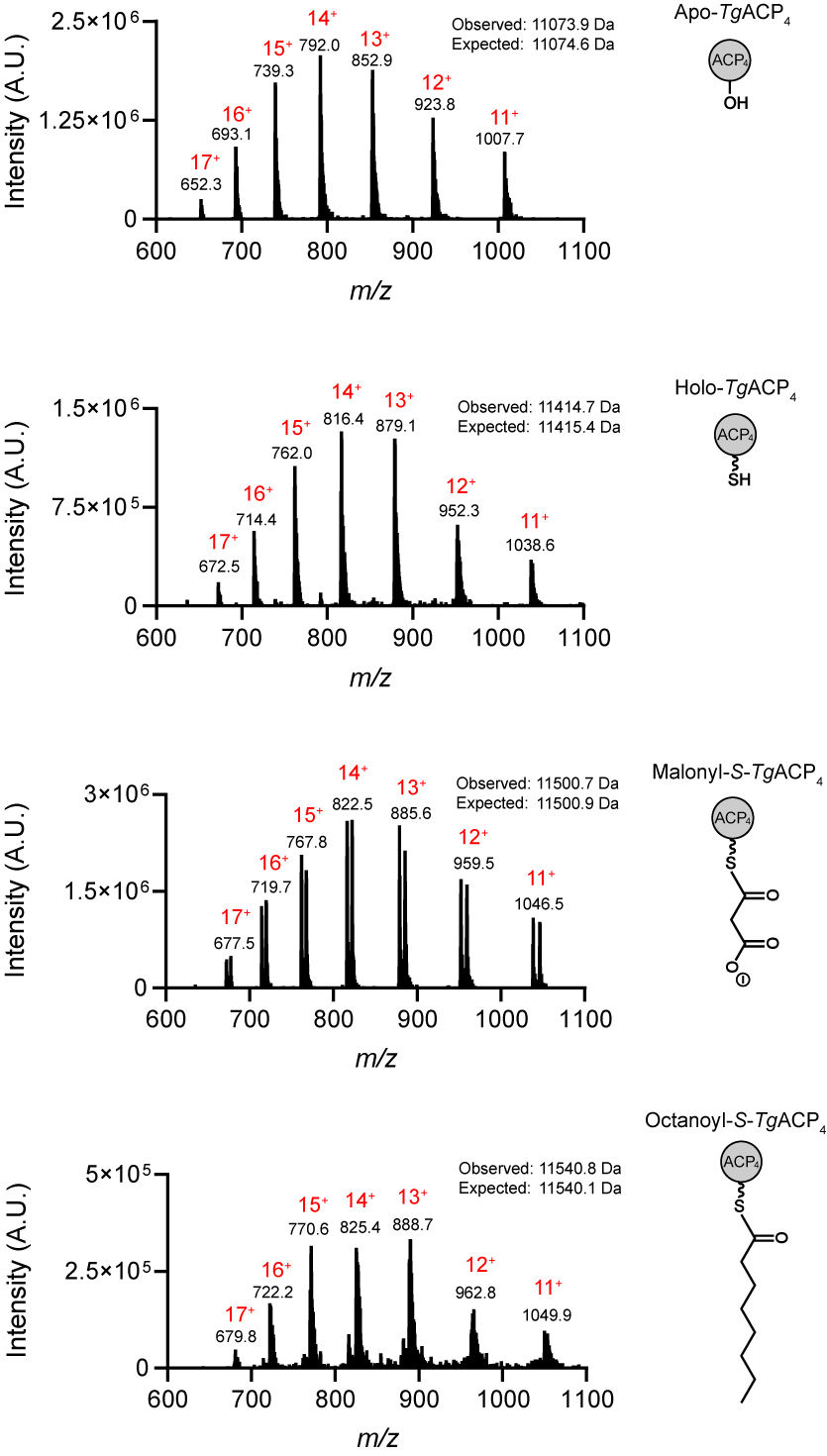

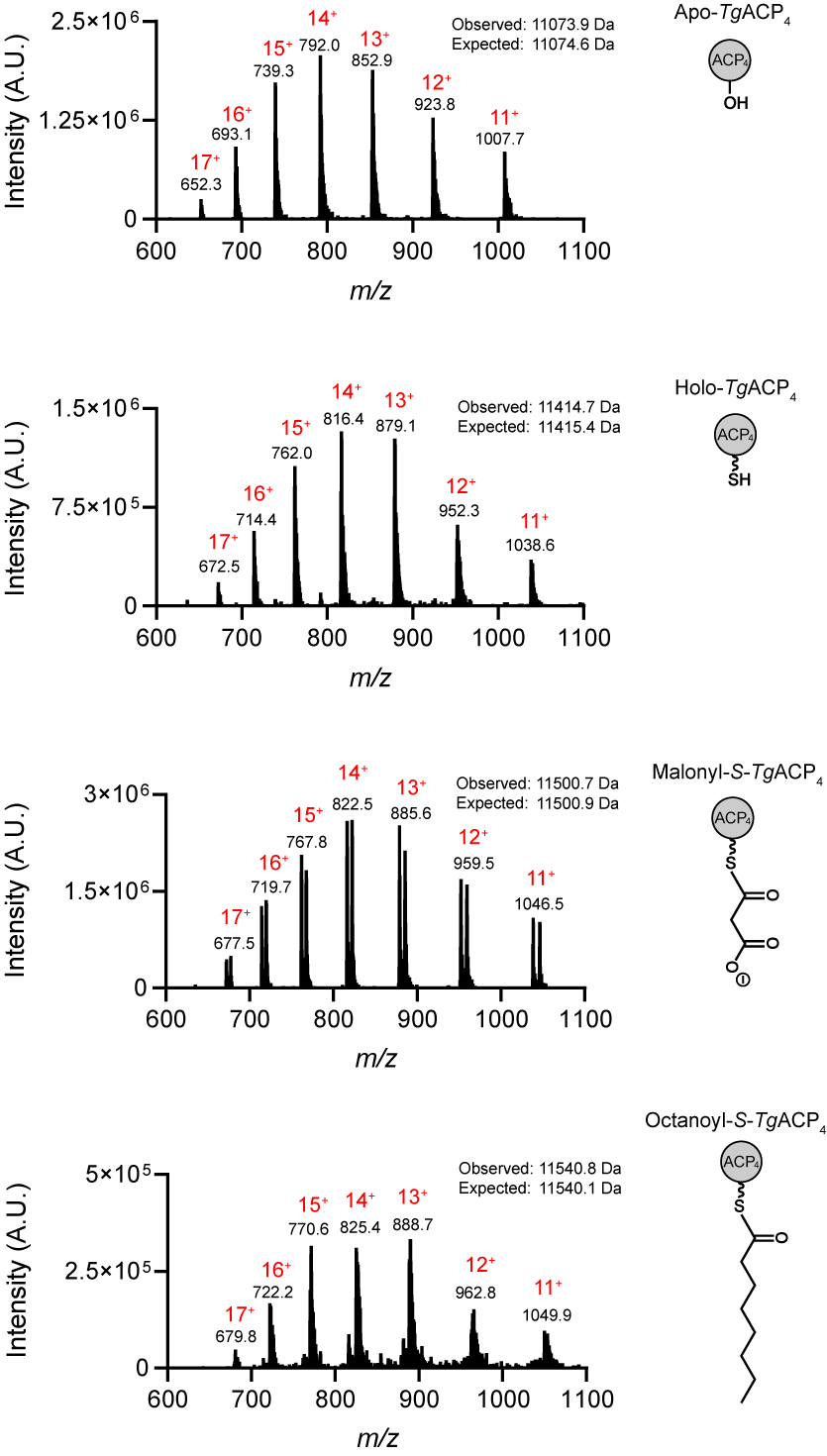

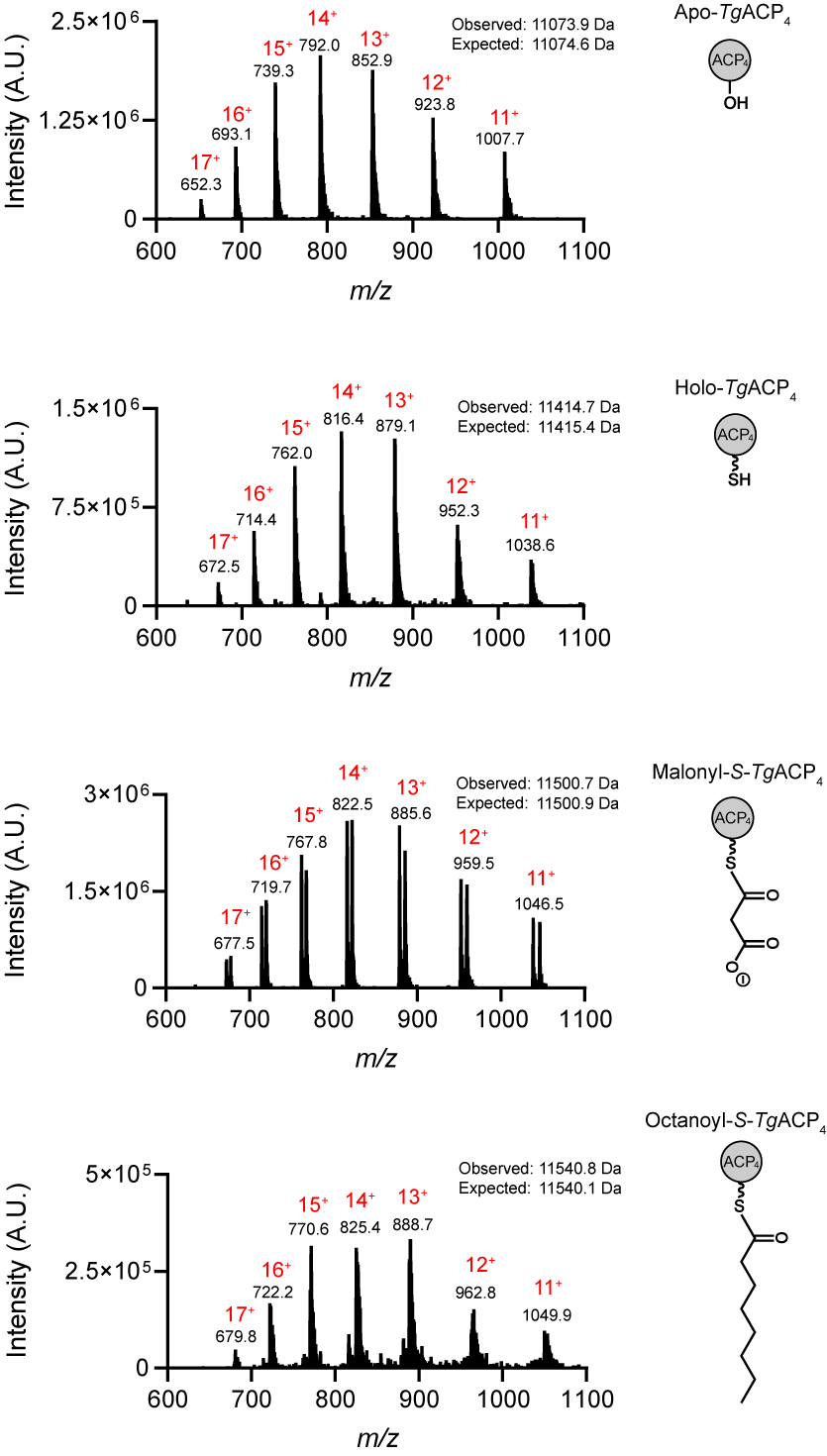

**D**

**C**

**Figure S6.** Analysis of ACP-derivatives by LCMS. Mass spectra of (A) apo-*Tg*ACP4, (B) holo-*Tg*ACP4, (C) malonyl-*S*-*Tg*ACP4, and (D) octanoyl-*S*-*Tg*ACP4 formed by incubation with Sfp and CoA substrate. Charge states and mass for each peak as well as observed and expected molecule weights are shown.

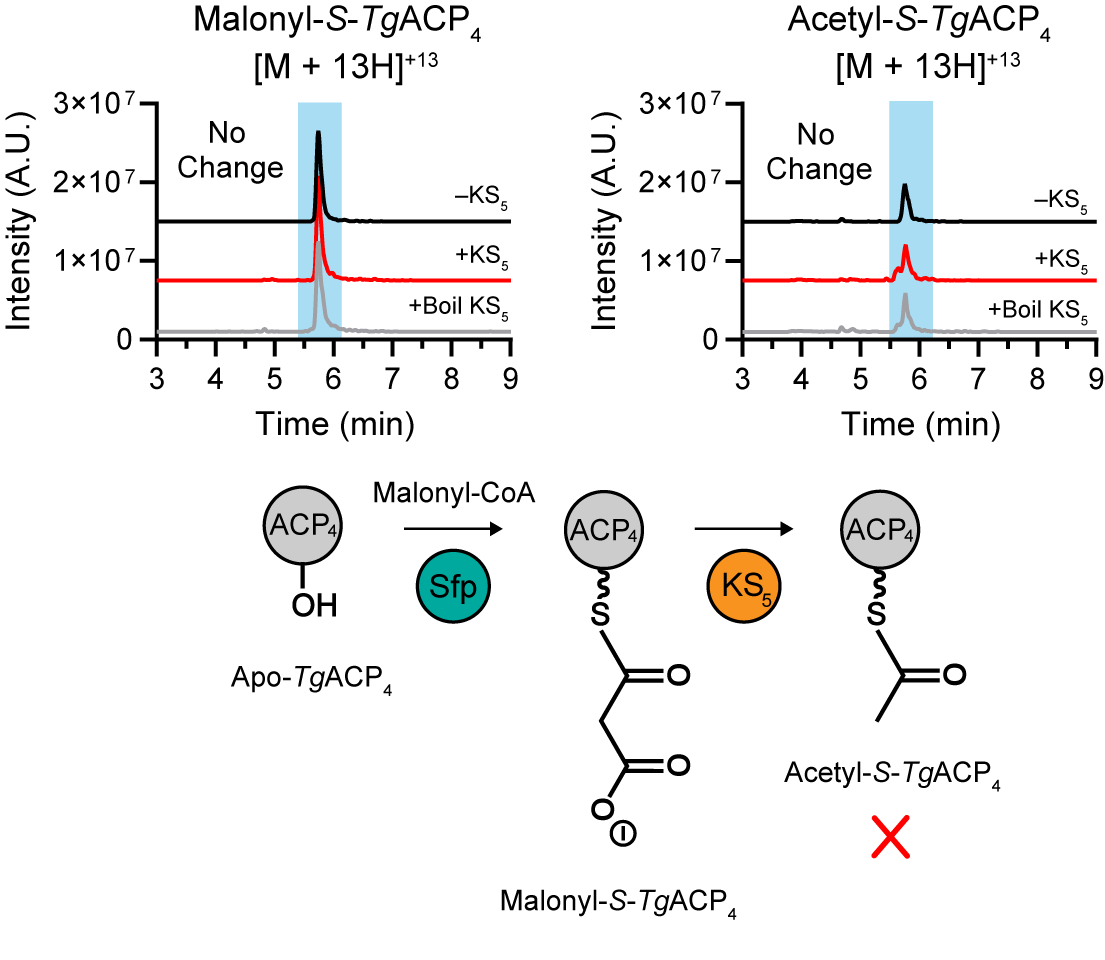

**Figure S7.** *In vitro* investigation of *Tg*KS5 domain. Representative LCMS extracted ion chromatograms of 13^+^ charge state for malonyl-*S*-*Tg*ACP4 and acetyl-*Tg*ACP4 with (red trace) and without (black trace) incubation with *Tg*TR. Reactions with boiled *Tg*TR (grey trace) included as a control. No increased formation of acetyl-*Tg*ACP4 was observed. Reaction scheme shown below plots.

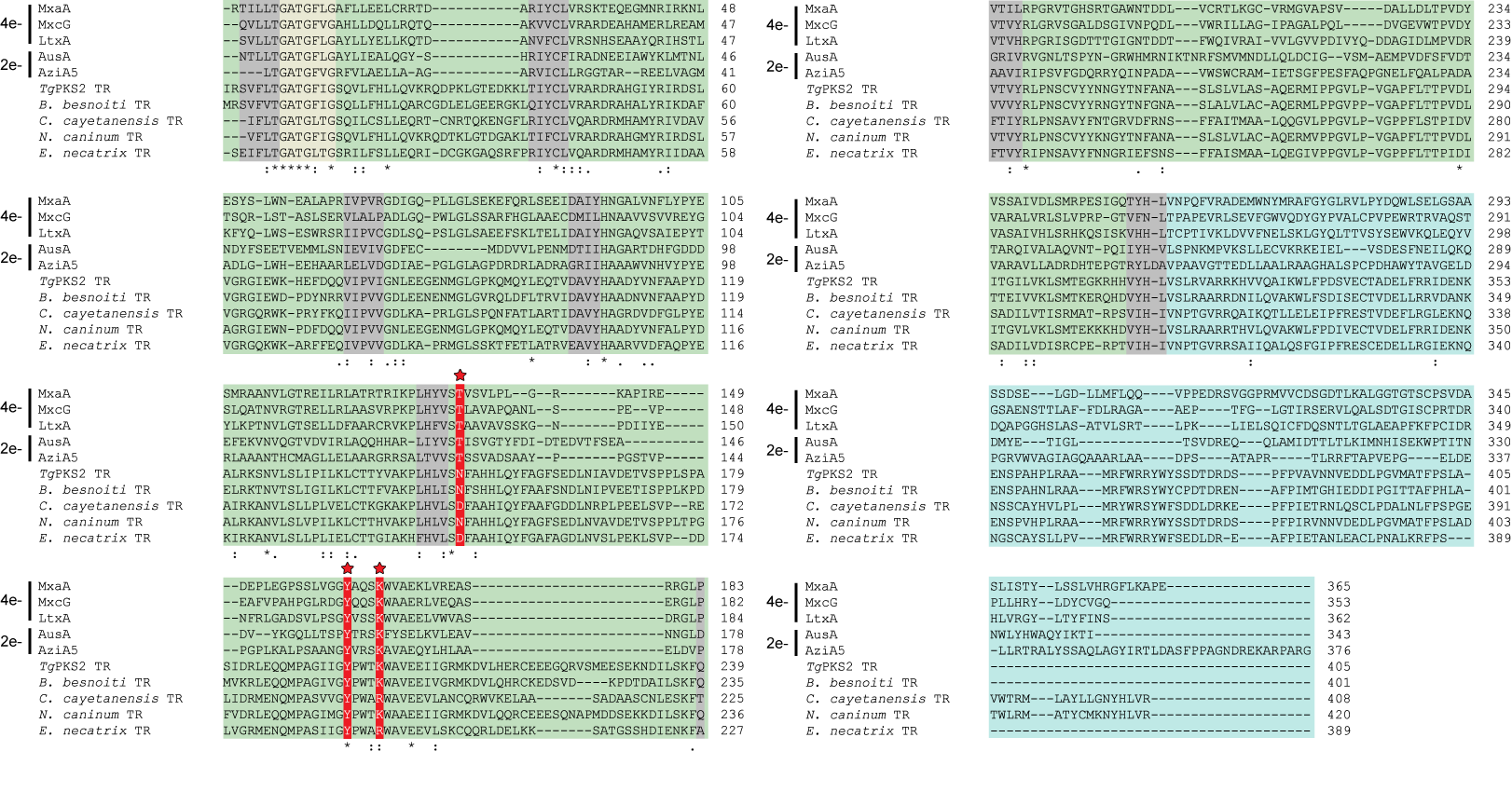
**Figure S8.** TR sequence alignments. ClustalW sequence alignments of *Tg*PKS2 TR, representative previously characterized 4e^-^ and 2e^-^ TR domains, and representative putative Coccidia TR domains. The predicted N-terminal domain (green) and C-terminal domain (blue) are highlighted and the predicted Rossman fold is highlighted (grey). The GxxGxxG motif implicated in cofactor binding is highlighted (tan). The catalytic triad residues are highlighted (red) and denoted by a red star. “*” denotes a residue that is identical in all sequences, “:” denotes residue substitutions with strongly similar properties within each sequence, and “.” denotes residue substitutions with weakly similar properties.

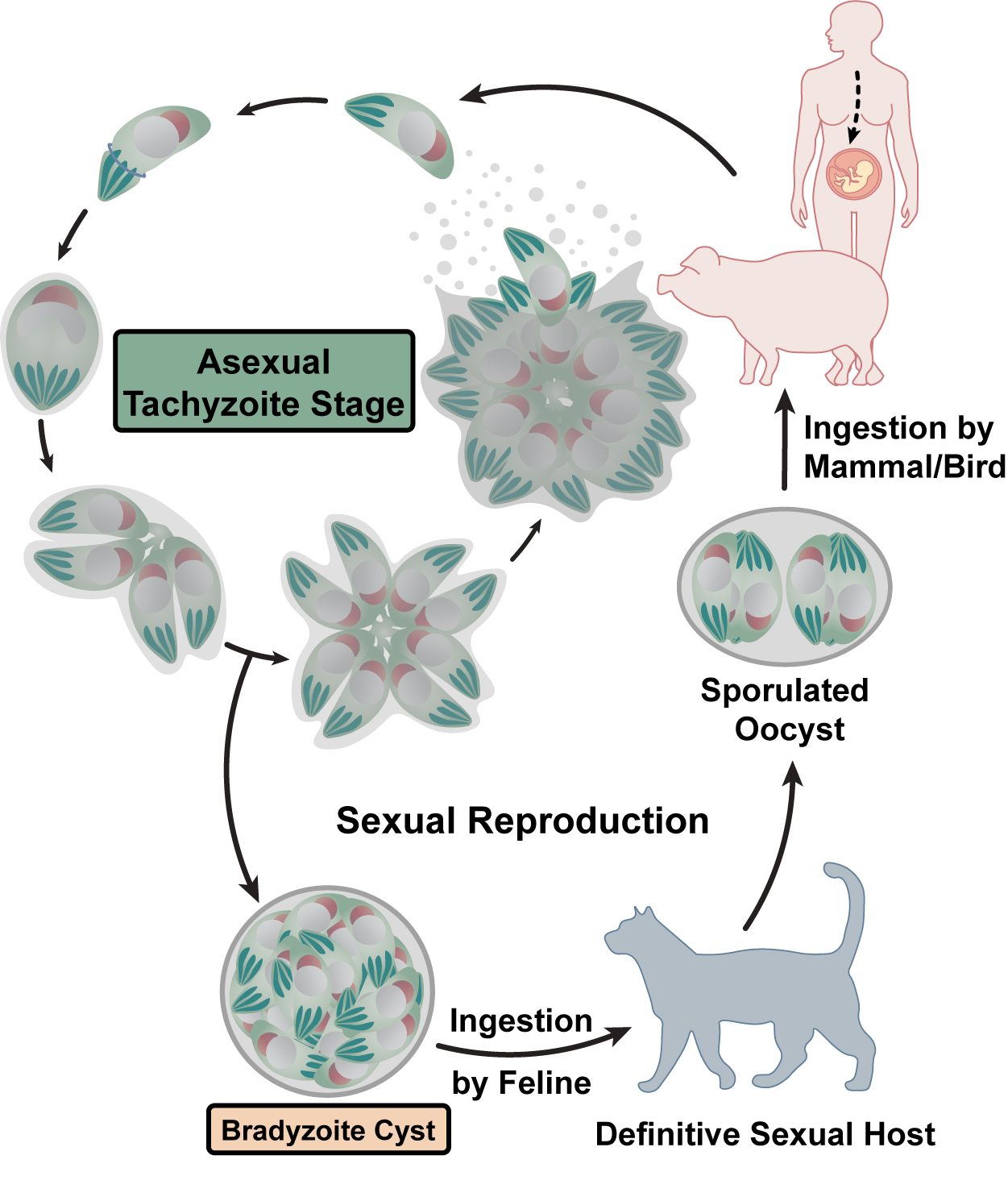

**Figure S9.** Overview of the *T. gondii* lifecycle. Sexual reproduction occurs in the definitive feline host, where oocysts are shed, sporulated, and can be up taken by virtually any warm-blooded host. *T. gondii* undergoes asexual reproduction during acute infection as tachyzoites and can differentiate into latent lipid-walled bradyzoite tissue cysts. These encysted bradyzoites can reactivate to relapse acute infection or can be taken up by the definitive sexual host, completing the lifecycle.
